# Fungal chromatin mapping identifies BasR, as the regulatory node of bacteria-induced fungal secondary metabolism

**DOI:** 10.1101/211979

**Authors:** Juliane Fischer, Sebastian Y. Müller, Tina Netzker, Nils Jäger, Agnieszka Gacek-Matthews, Kirstin Scherlach, Maria C. Stroe, María García-Altares, Francesco Pezzini, Hanno Schoeler, Michael Reichelt, Jonathan Gershenzon, Mario K. C. Krespach, Ekaterina Shelest, Volker Schroeckh, Vito Valiante, Thorsten Heinzel, Christian Hertweck, Joseph Strauss, Axel A. Brakhage

**Author notes:** equal contribution (Juliane Fischer, Sebastian Y. Müller, Tina Netzker, Nils Jäger). current address: Department of Plant Sciences, University of Cambridge, Downing Street, Cambridge CB2 3EA, UK. current address: Bioinformatics Unit, German Centre for Integrative Biodiversity Research (iDiv), Deutscher Platz 5e, 04103 Leipzig, Germany (Francesco Pezzini, Ekaterina Shelest). corresponding authors: Axel A. Brakhage, Joseph Strauss.

## Abstract

The eukaryotic epigenetic machinery is targeted by bacteria to reprogram the response of eukaryotes during their interaction with microorganisms. In line, we discovered that the bacterium *Streptomyces rapamycinicus* triggered increased chromatin acetylation and thus activation of the silent secondary metabolism *ors* gene cluster leading to the production of orsellinic acid in the fungus *Aspergillus nidulans*. Using this model we aim at understanding molecular mechanisms of communication between bacteria and eukaryotic microorganisms based on bacteria-triggered chromatin modification. By genome-wide ChIP-seq analysis of acetylated histone H3 (H3K9ac, H3K14ac) we uncovered the unique chromatin landscape in *A. nidulans* upon co-cultivation with *S. rapamycinicus*. Genome-wide acetylation of H3K9 correlated with increased gene expression, whereas H3K14 appears to function in transcriptional initiation by providing a docking side for regulatory proteins. In total, histones belonging to six secondary metabolism gene clusters showed higher acetylation during co-cultivation including the *ors*, aspercryptin, cichorine, sterigmatocystin, anthrone and 2,4-dihydroxy-3-methyl-6-(2-oxopropyl)benzaldehyde gene cluster with the emericellamide cluster being the only one with reduced acetylation and expression. Differentially acetylated histones were also detected in genes involved in amino acid and nitrogen metabolism, signaling, and genes encoding transcription factors. In conjunction with LC-MS/MS and MALDI-MS imaging, molecular analyses revealed the cross-pathway control and Myb-like transcription factor BasR as regulatory nodes for transduction of the bacterial signal in the fungus. The presence of *basR* in other fungal species allowed forecasting the inducibility of ors-like gene clusters by *S. rapamycinicus* in these fungi, and thus their effective interaction with activation of otherwise silent gene clusters.

## Introduction

The eukaryotic epigenetic machinery has been shown to be targeted by bacteria. For example, bacteria can secrete chromatin modifiers or proteins such as methyltransferases that silence chromatin of eukaryotic cells (1, 2). As an early example, we discovered that the silent secondary metabolite (SM) gene cluster for orsellinic acid (*ors*) in the filamentous fungus *Aspergillus nidulans* is activated upon physical interaction with the bacterium *Streptomyces rapamycinicus*. The interaction of the fungus with this distinct bacterium led to increased acetylation of histone H3 lysine 9 and 14 at the *ors* gene cluster and thus to its activation (3–5). The lysine acetyltransferase (KAT) responsible for the acetylation and activation of the *ors* gene cluster was shown to be GcnE (4).

Using this model we aim at understanding the molecular mechanisms of microbial communication based on bacteria-triggered chromatin modification. In order to obtain a holistic view on the fungal bacterial interaction that in the future might allow predicting interaction partners and discovering the molecular elements involved, we developed a genome-wide chromatin immunoprecipitation (ChIP)-seq analysis specifically during co-cultivation. This led to the discovery of major alterations of epigenetic marks in the fungus triggered by the bacterium and the identification of BasR as the key regulatory node required for linking bacterial signals with the regulation of SM gene clusters.

## Results

### Genome-wide profiles of H3K9 and H3K14 acetylation in *A. nidulans* change upon co-cultivation with *S. rapamycinicus*

*A. nidulans* with and without *S. rapamycinicus* was analyzed by genome-wide ChiP-seq for enrichment of acetylated (ac) histone H3 at lysines K9 and K14 (Fig. 1; SI Results-Details of ChIP analysis). To account for reads originating from *S. rapamycinicus* we fused the genomes of *A. nidulans* (8 chromosomes) and *S. rapamycinicus*. The resulting fused genome also served as reference for mapping of chromatin marks (see SI Results – Details of the ChIP analysis).

**Figure 1.**
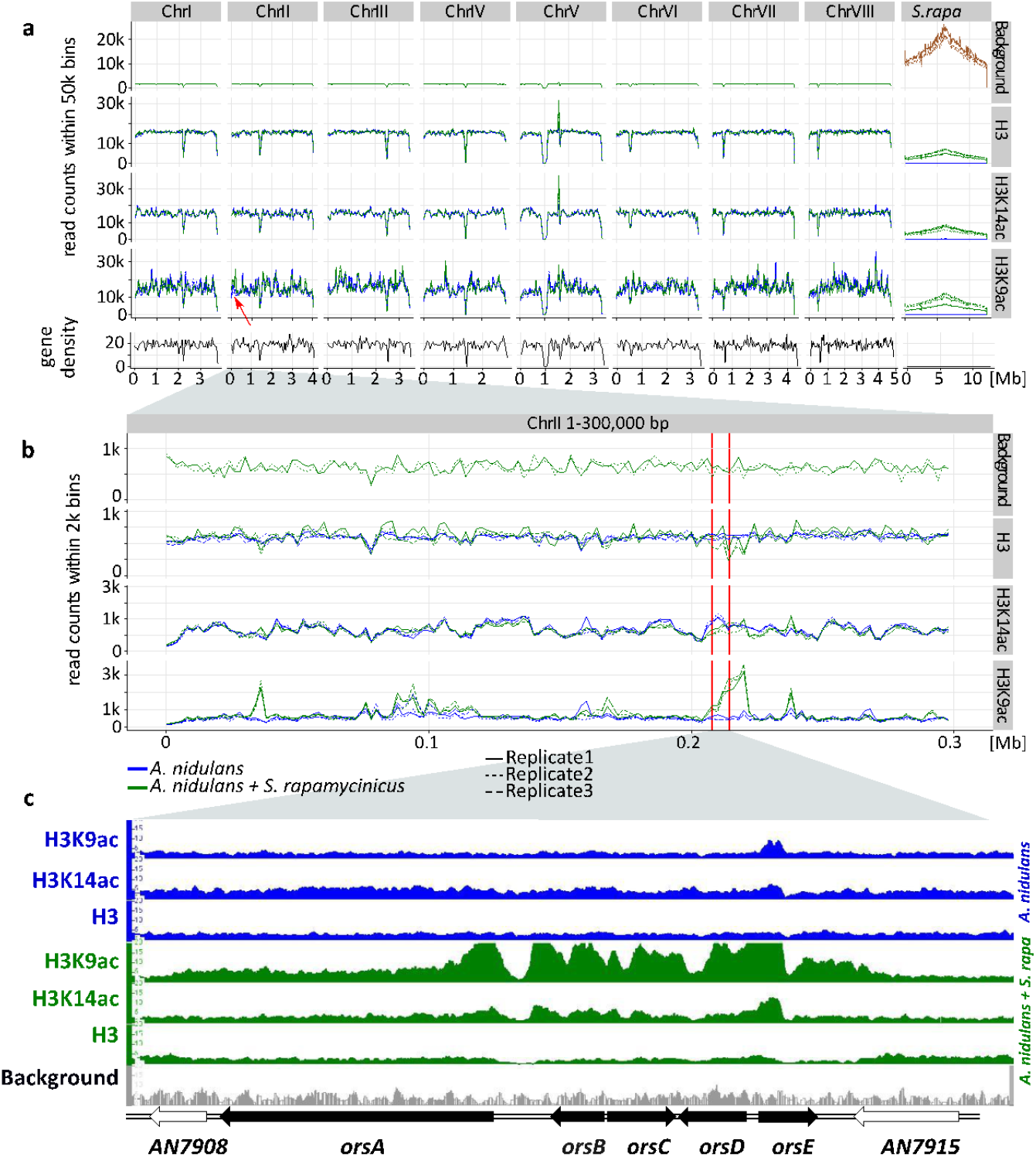
Genome-wide coverage plot of the fused fungal-bacterial genome with indication of H3(Cterm) and acetylated H3 (K9 and K14). For each condition, ChIP-seq analyses of three independent samples were performed. (**a**) Genome-wide analysis covering all chromosomes. Data for all the chromosomes I to VIII of *A. nidulans* as well as for the chromosome of *S. rapamycinicus* are shown. X axis corresponds to genome coordinates of the fused genome in Mb. Y axis corresponds to the number of reads mapping within equally sized windows (bins) which segment the fused genome at a resolution of 50 kb for each library separately (see methods for details). The read count values are plotted at the midpoint of each bin which then were connected by lines. Gene density is reported likewise by counting the number of genes for each bin instead of reads. Background values derive from *S. rapamycinicus* (brown) and *A. nidulans* (green) grown in monoculture. The red arrow indicates the location of the *ors* gene cluster. (**b**) Zoom into chromosome II. The red lines mark the *ors* gene cluster. Data of three replicates are shown, which show the same tendency. Overall intensity of background, H3K9ac, H3K14ac and H3(Cterm) compared between *A. nidulans* monoculture (blue) and co-culture (green) is shown, as well as the average genome density (black). (**c**) Example of an IGV screenshot showing the region of the *ors* gene cluster at the bottom of the figure labeled with black arrows. Other differentially acetylated gene bodies are listed in table 1. Blank gene arrows indicate genes not belonging to the *ors* gene cluster. Data obtained from monocultures of the fungus are depicted in blue, from co-cultivation in green and background data in grey.

H3K14ac and H3K9ac showed a higher degree of variability across the genome compared to H3 implying a more specific regulatory dynamics by histone acetylation than through H3 localization. Some areas such as a region in the first half on chromosome 4 were particularly enriched in those marks, potentially marking distinctive chromatin domains. A domain particularly enriched for H3K9ac was found around the *ors* gene cluster (Figs. 1 c & 2), thus supporting our previous data (4). The coverage profiles of H3, H3K14ac and H3K9ac have consistently changed in co-culture compared to monoculture as seen in Fig 1. In particular the promoter region of the genes *orsD* and *orsA* showed reduced nucleosome occupancy (see Fig. 1 c). This could be due to a redistribution of nucleosomes which ultimately changes the distribution of histone marks. The changes of H3K14ac in Fig. 1 are therefore likely due to nucleosome rearrangements towards the translation start sites (TSS) rather than increased amounts of this modification. This is supported by the observation that unmodified H3 was strongly depleted throughout the *ors* cluster, especially at the *orsA* and *orsD* TSS.

**Figure 2.**
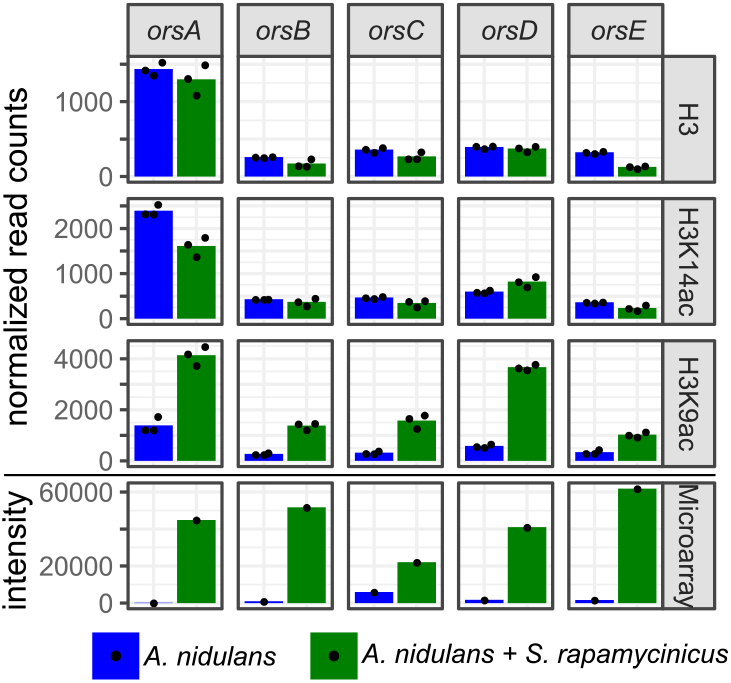
Normalized read counts derived from differential chromatin state (DCS) analysis obtained for the *ors* genes based on H3, H3K14ac and H3K9ac ChIP-seq. Data were generated for the area 500 bp down- and 1000 bp upstream from the TSS. Depicted bars are calculated from three data points.

### Co-cultivation of *A. nidulans* with *S. rapamycinicus* had a major impact on SM gene clusters, nitrogen assimilation, signalling and mitochondrial activity

We employed two strategies to determine changes of histone modification levels. The first analysis was based on the finding that histone acetylation can mostly be found on histones within a gene, in particular on nucleosomes +1 and +2 (6) (Fig. S1 a). We therefore counted mapped reads overlapping genes for each library. This formed the basis for a quantitative comparison between monocultures and co-cultures using standard read counting methods for sequencing data (see methods). Throughout this study, we refer to this method as differential chromatin state (DCS) analysis. The second analysis was based on a first round of peak-calling and subsequent quantification of the peaks. Comparison of the generated data sets showed 84 ± 1.7 % similarity. The data obtained from the gene-based DCS method (Table S1) were used for both further analyses and comparison of the culture conditions using a false discovery rate (FDR) cut-off of 0.01. This does not include further filtering on the log-fold changes (LFCs) to capture possible biological relevance of the detected changes. Quality and absence of possible biases introduced by the co-culture or other sources were further investigated by MA plots. They showed a symmetrical and even distribution around LFC = 0, meeting the requirements for the statistical tests described in methods (Fig. S2). DCS analysis of H3, as a proxy for nucleosome occupancy, was found to be lower (FDR < 0.01) in 37 genes and higher in 2 genes during co-cultivation. Using the same cut-off, during bacterial-fungal co-cultivation H3K14ac levels were found to be lower for 154 genes and higher for 104 genes. Differential acetylation of chromatin was found for H3K9ac with 297 genes with lower and 593 with significantly higher acetylation (Table S1).

The analysis of microarray data obtained under identical conditions showed a positive correlation of higher gene expression with H3K9 acetylation (r=0.2 for all genes and r=0.5 for a subset of genes showing differential acetylation; Figs. S3 & S4). Data for selected genes are summarized in Table S2 showing the LFCs of H3K9ac ChIP-seq data with their corresponding microarray data. In total, histones belonging to six SM gene clusters showed higher acetylation during co-cultivation including the *ors*, aspercryptin, cichorine, sterigmatocystin (stc), anthrone (*mdp*) and 2,4-dihydroxy-3-methyl-6-(2-oxopropyl)benzaldehyde (DHMBA) gene cluster with the emericellamide *(eas)* cluster being the only one with reduced acetylation and expression (Table S2, section V). With a few exceptions genes covered by histone H3 with higher acetylation are involved in calcium signaling and asexual development (Table S2, sections III & IV; Fig S5).

A major group of genes with lower acetylation in mixed cultivation compared to the monoculture of *A. nidulans* is linked to the fungal nitrogen metabolism (Table S2, section I) including genes for the utilization of primary and secondary nitrogen sources such as genes of the nitrate assimilation gene cluster and the glutamine dehydrogenase gene (Figs. S6 & 3 a). These data were confirmed by quantifying the expression of identified genes by qRT-PCR (Fig. 3 b).

**Figure 3.**
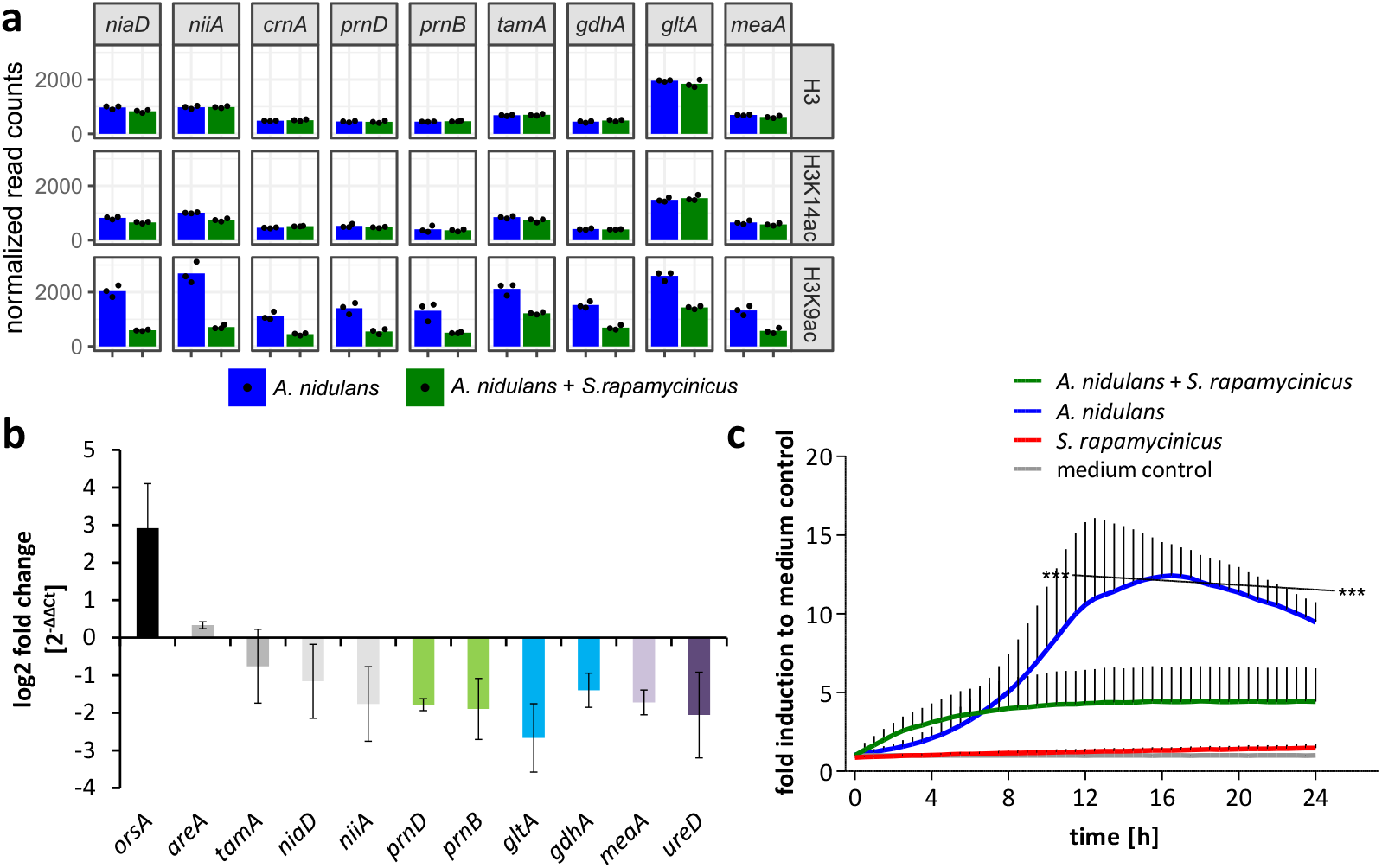
Influence of *S. rapamycinicus* on the fungal nitrogen metabolism and mitochondrial functions. (**a**) Normalized ChIP-seq read counts were used to quantify chromatin state (H3, H3K14ac, H3K9ac) for nitrogen metabolism genes. Counts were obtained by counting reads mapping to the promoter area for each gene that is 500 bp down- and 1000 bp upstream from the TSS. Depicted bars are calculated from three data points. (**b**) Transcription analysis of randomly selected genes of primary and secondary nitrogen metabolism by qRT-PCR during co-cultivation. Relative mRNA levels were measured after 3 hours and normalized to the β-actin gene expression. The transcription of *orsA* was used as a positive control. (**c**) Respiratory activity comparing *A. nidulans* grown in co-culture with *S. rapamycinicus* and *A. nidulans* in monoculture. Respiratory activity was determined using a resazurin assay. Data were normalized to medium. The black line shows the time points that are significantly different between *A. nidulans* and *A. nidulans* grown in co-culture with *S. rapamycinicus*. *** p < 0.001

Genes assigned to mitochondrial function showed decreased acetylation of H3K9 which implied reduced mitochondrial function. This assumption was confirmed by measuring the respiratory activity of fungal cells. In monoculture the fungus showed a high metabolic activity, which was significantly reduced during co-cultivation (Fig. 3 c).

### Bacteria induce elements of the fungal cross-pathway control

To identify transcription factors involved in transducing the bacterial signal to the fungal expression machinery and because a transcription factor gene is missing in the *ors* gene cluster, we analyzed the 890 differentially H3K9 acetylated genes for those annotated as putatively involved in transcriptional regulation. 22 putative transcription factor-encoding genes fulfilled this requirement (Table S2, section VII). Most of them (18 genes) showed significantly higher acetylation in co-culture, while only 4 genes had lower acetylation. Among the higher acetylated genes were *cpcA*, coding for the central transcriptional activator of the cross-pathway control CpcA, as well as the bZIP transcription factor gene *jlbA* (jun-like bZIP). Both genes have been shown to be highly expressed during amino acid starvation in *A. nidulans* (7, 8). Additionally, a putative orthologue *(AN7174)* of the *S. cerevisiae bas1* gene showed an increased acetylation. In yeast, together with the homeodomain protein Bas2p, Bas1p is involved in the regulation of amino acid biosynthesis (9, 10). Consistently, a number of genes related to amino acid metabolism showed increased acetylation for H3K9 during the co-cultivation of *A. nidulans* with *S. rapamycinicus* (Table S2, section II). To correlate the ChIP-seq data with expression levels of *cpcA, jlbA* and *AN7174* qRT-PCR analysis was carried out, which demonstrated up-regulation of *cpcA* and *AN7174* during co-cultivation (Fig. 4 a). In *S. cerevisiae* it was shown that Gcn4 (CpcA in *A. nidulans)* and Bas1p share a similar DNA-binding motif and both activate the transcription of the histidine biosynthesis gene *HIS7* independently from each other (9). In line of a possible involvement of these transcription factors is the observation that the addition of the histidine analogue 3-aminotriazole (3-AT), which is known to induce the cross-pathway control (CPC) *via* amino acid starvation, led to the production of orsellinic acid in the fungal monoculture (Fig. 4 b) and to an increased expression of *orsA, cpcA* and *AN7174* (Fig. 4 c).

**Figure 4.**
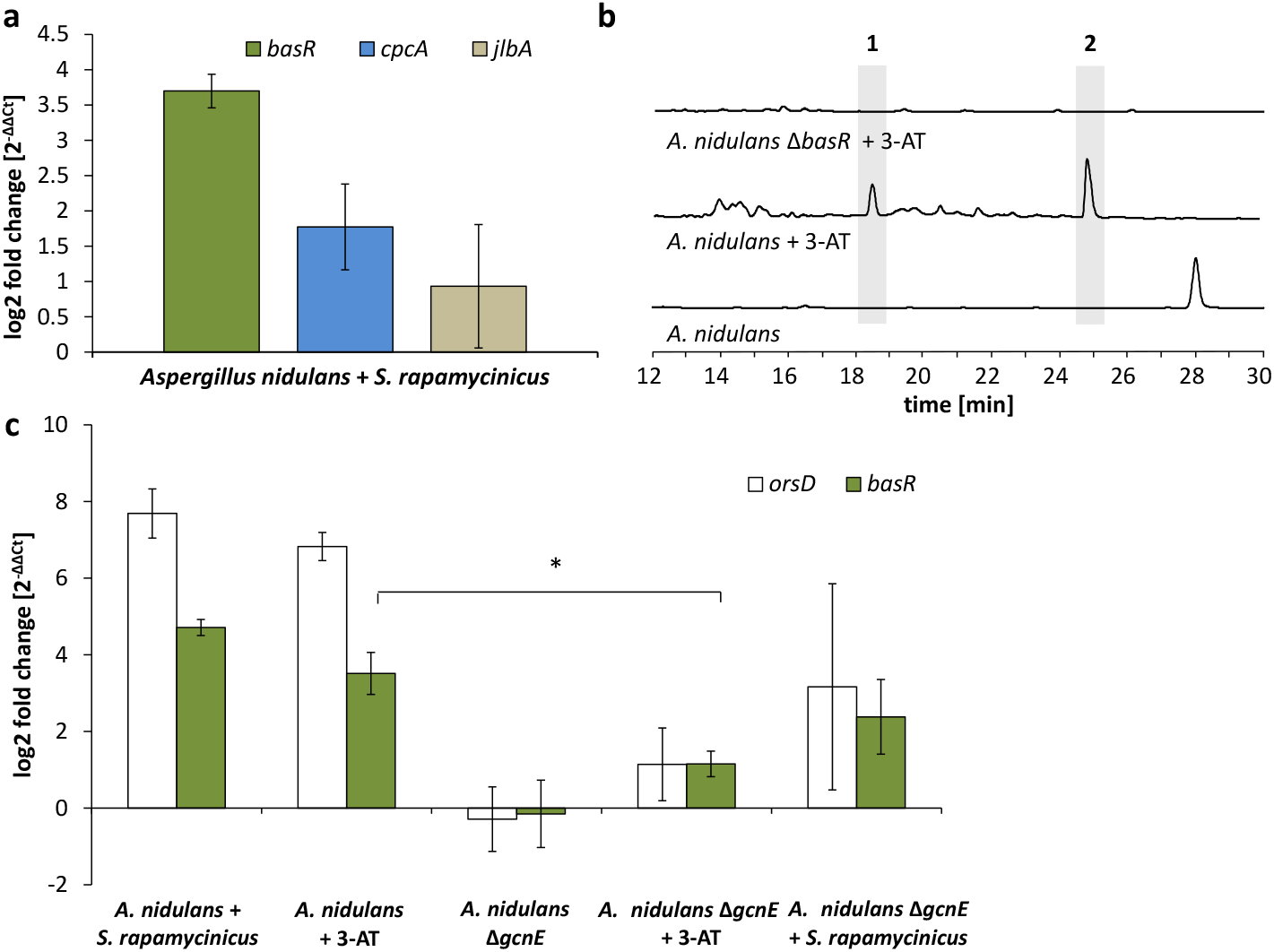
Artificial histidine starvation using 3-AT led to *ors* gene cluster activation. (**a**) Transcription of *basR, cpcA* and *jlbA* determined by qRT-PCR after 3h of co-cultivation. Relative mRNA levels were compared to β-actin gene expression. (**b**) HPLC-based detection of orsellinic acid (1) and lecanoric acid (2) in supernatants of *A. nidulans* cultures treated with 3-AT. (**c**) Relative transcript levels of *orsA, cpcA* and *basR* 6 hours after 3-AT addition to the *A. nidulans* monoculture and the *gcnE* deletion mutant. *p < 0.05

To analyze a possible involvement of these genes in the bacteria-induced activation of the *ors* gene cluster, the genes *cpcA* (data not shown) and *AN7174* gene (Fig. S7 a) were deleted. Deletion of *cpcA* in *A. nidulans* showed no effect on the induction of the *ors* gene cluster in response to *S. rapamycinicus* (data not shown), while deletion of *AN7174* resulted in a significantly reduced expression of *orsA* and *orsD*, and in complete loss of orsellinic acid production (Fig. 5). Therefore, *AN7174* was named *basR* and analyzed in detail.

**Figure 5.**
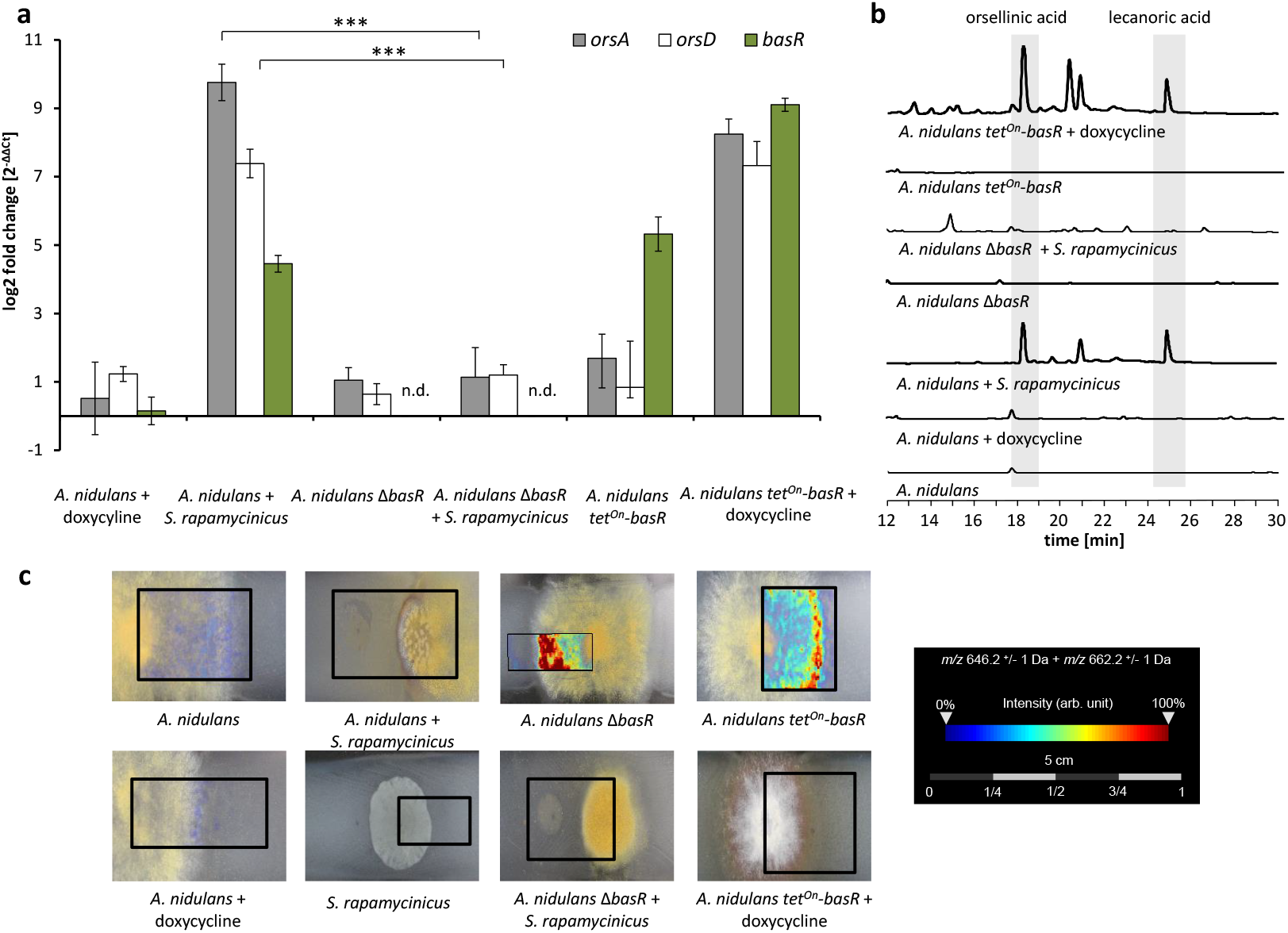
The Myb-like transcription factor BasR of *A. nidulans* is required for *S. rapamycinicus-triggered* regulation of SMs. (**a**) Relative transcript levels of *ors* cluster genes *orsA, orsD* and *basR* after 6 hours of cultivation in Δ*basR* mutant strain and *tet*^on^-*basR* overexpression strain incubated with and without doxycycline. Transcript levels were measured by qRT-PCR normalized to β-actin transcript levels. (**b**) HPLC-based detection of orsellinic and lecanoric acid in wild-type strain, *basR* deletion mutant and *basR* overexpression strain. (**c**) Visualization of ions *m/z* 646.3 and *m/z* 662.3 +/− 1 Da, potentially corresponding to [M+Na]^+^ and [M+K]^+^ of emericellamide E/F (C_32_H_57_N_5_O_7_; accurate mass 623.4258), by MALDI-MS imaging. Images corrected by median normalization and weak denoising. n.d.: not detectable; ***p < 0.001

### The transcription factor BasR is the central regulatory node of bacteria-triggered SM gene cluster regulation

Further analysis of the *A. nidulans* genome revealed a second gene (*AN8377*) encoding a putative orthologue of the *S. cerevisiae bas1* gene (Figs. 6 & S8). Both genes (*basR* & *AN8377*) code for Myb-like transcription factors whose function in filamentous fungi is completely unknown. We compared the H3K9 acetylation and gene expression of both genes upon co-cultivation. The *basR* gene showed higher H3K9 acetylation (LFC = 0.6) and drastically increased transcription (LFC = 5.85) during co-cultivation compared to *AN8377* (H3K9ac LFC = −0.03; Microarray LFC = 0.14). Deletion of *AN8377* (Fig. S9 a) did not affect the induction of fungal orsellinic acid production upon co-cultivation (Fig. S9 b), excluding a role of *AN8377* in this process.

**Figure 6.**
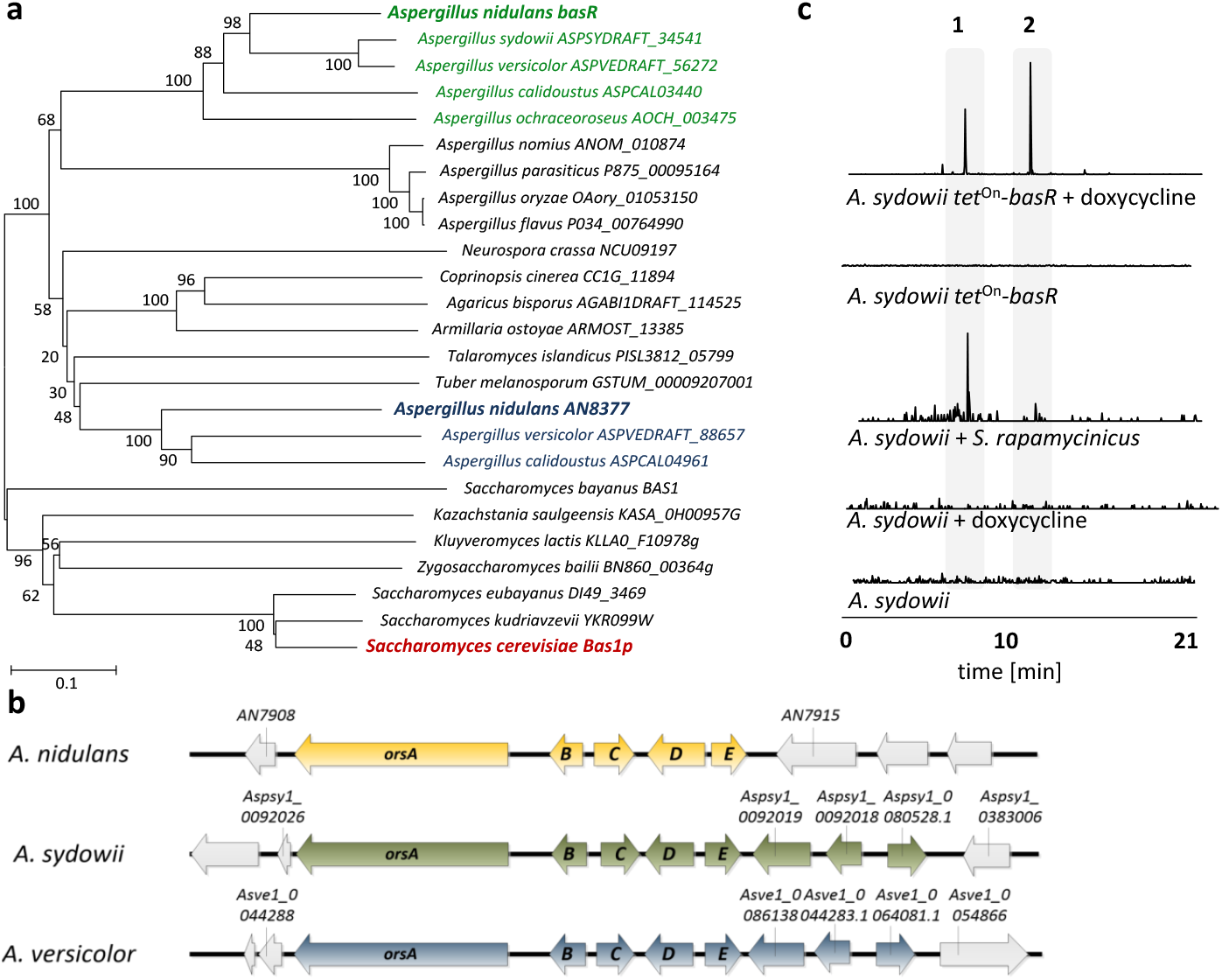
Co-occurrence of BasR and the orsellinic acid gene cluster in other fungi is linked to the *S. rapamycinicus* triggered *ors* gene cluster activation. (**a**) Phylogenetic analysis of BasR (*AN7174;* green) showing its position among other fungi. The percentage of trees in which the associated taxa clustered together is shown next to the branches. The names of the selected sequences are given according to their UniProt accession numbers. (**b**) Alignment of the orsellinic acid gene clusters in the fungal species containing a *basR* homologue (*A. nidulans, A. sydowii, A. versicolor*), where *orsA* encodes the polyketide synthase, while *orsB-orsE* code for tailoring enzymes. (**c**) LC-MS-based detection of orsellinic and lecanoric acid in monoculture of the *A. sydowii basR* overexpression strain following induction with doxycycline (left) and during co-cultivation of *A. sydowii* and *S. rapamycinicus* (right). LC-MS profiles of the extracted ion chromatogram (EIC) are shown for *m/z* 167 [M − H]^−^, which corresponds to orsellinate. Orsellinic (1) and lecanoric acid (2) were detected *via* its fragment ion orsellinate.

In *S. cerevisiae* Bas1p needs the interaction with Bas2p for the transcriptional activation of several genes required for histidine and purine biosynthesis (9). The C-terminal activation and regulatory (BIRD) domain of Bas1, which was described to mediate this Bas1p-Bas2p interaction (11), is missing in BasR. It is thus not surprising that in the *A. nidulans* genome we did not find an orthologue for the *S. cerevisiae bas2* gene. While the addition of 3-AT to monocultures of *A. nidulans* led to the production of orsellinic acid and derivatives thereof, the effect of 3-AT was abolished in the *basR* deletion mutant strain (Fig. 4 b).

As the transcriptional activation of *HIS7* by Bas1/Bas2 upon adenine limitation in yeast requires a functional Gcn5 (GcnE in *A. nidulans*) (10), we raised the question whether GcnE is needed for full *basR* expression. Addition of *S. rapamycinicus* or 3-AT to the *gcnE* deletion mutant led to decreased *basR* gene expression compared to the co-culture or a monoculture of the wild type with 3-AT (Fig. 4 c). These data indicate that GcnE is required for *basR* expression. Inspection of the *basR* mutant strain on agar plates did not reveal further obvious phenotypes (data not shown).

To further substantiate the influence of *basR* on the *ors* gene cluster we generated a *basR* overexpression strain (Fig. S7 b) by employing the inducible *tet*^On^ system (12). Addition of doxycyline to the media induced *basR* expression as well as the expression of the *ors* gene cluster (Fig. 5 a). However, *basR* gene expression was already detectable without doxycycline addition, indicating “leakiness” of the *tet*^On^ system. Nevertheless, production of orsellinic and lecanoric acid was only detected upon doxycycline addition (Fig. 5 b), supporting the important role of BasR for their biosynthesis. To address the question whether other SM biosyntheses are regulated by BasR we searched for other metabolites. Obvious candidates were the emericellamides, as the acetylation of the corresponding gene cluster was decreased during co-cultivation (Table S2). This finding was perfectly mirrored when we applied MALDI-mass spectrometry (MS) imaging which showed reduced levels of emericellamides in both *basR*-overproducing colonies of *A. nidulans* and in co-grown colonies in contrast to colonies without the streptomycete (Fig. 5 c).

### The presence of BasR in fungal species allows forecasting the inducibility of ors-like gene clusters by *S. rapamycinicus*

To address the question whether *basR* homologues exist in other fungi and whether potential homologues have similar functions, we analyzed fungal genomes using BlastP. Surprisingly, obvious *basR* homologues are only present in few other *Aspergillus* spp. including *Aspergillus sydowii* and *Aspergillus versicolor*, but are apparently lacking in many others (Fig. S8). Interestingly, except for three additional genes in both fungi a gene cluster similar to the *ors* gene cluster of *A. nidulans* was also identified (Fig. 6 a). We overexpressed *basR* in *A. sydowii* using the *tet*^On^ system to analyze its function (Fig. S10). LC-MS analyses revealed the appearance of novel masses which were assigned to orsellinic acid derivatives (Figure 6 b).

Finally, we addressed the question whether the presence of the *basR* gene and the *ors* gene cluster allows forecasting their inducibility by *S. rapamycinicus*. As shown in Fig. 7, also co-cultivation of *A. sydowii* with *S. rapamcinicus* led to the activation of the fungal *ors* gene cluster, again linking BasR with bacteria-triggered induction of the production of orsellinic acid derivatives.

**Figure 7.**
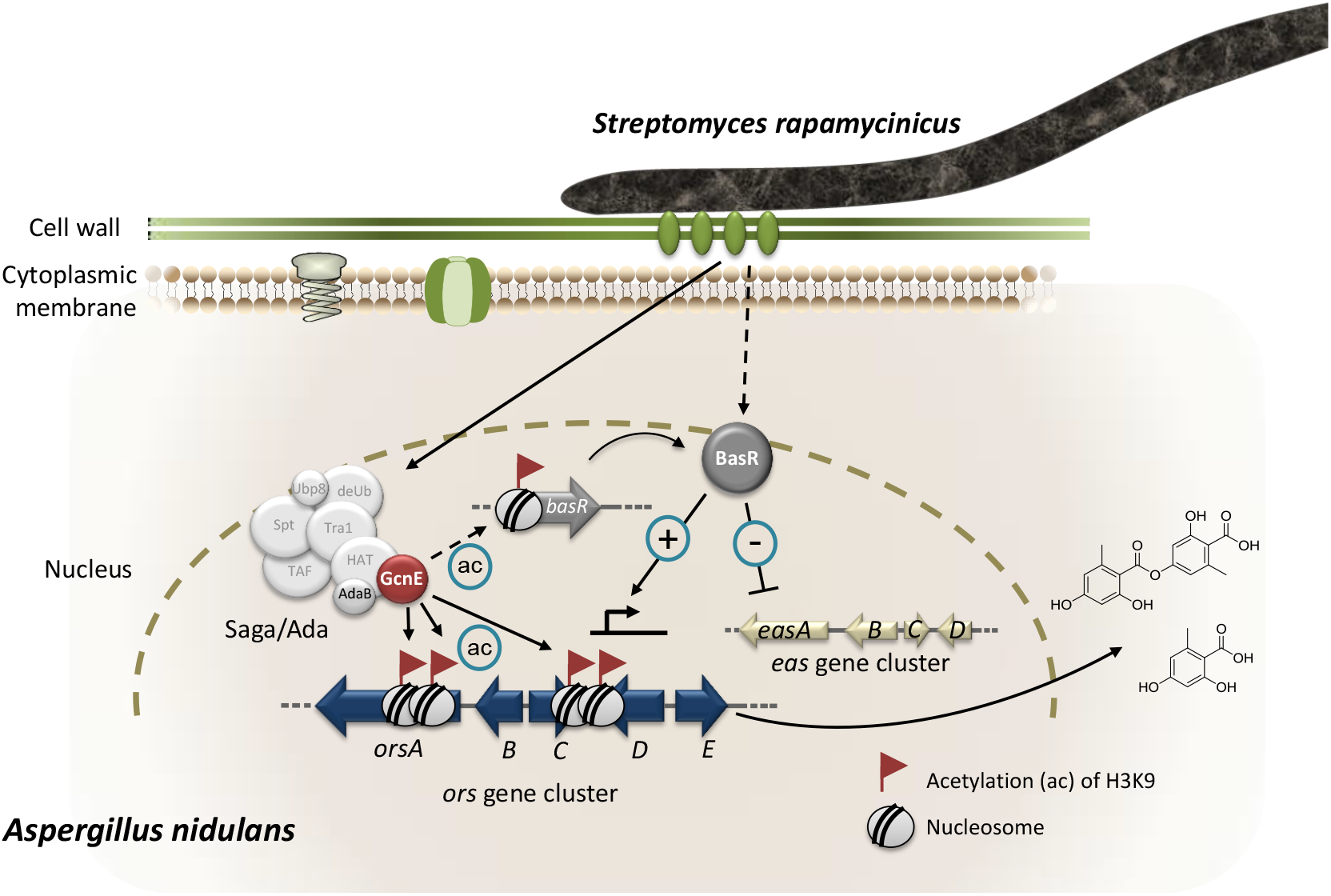
Model of the *S. rapamycinicus – A. nidulans* interaction. Co-cultivation leads to activation of the *basR* gene. The lysine acetyltransferase GcnE specifically acetylates (ac) lysine (K) 9 of histone H3 at the *ors* gene cluster and presumably at the *basR* gene promoter. As a consequence, *basR* is expressed. The transcription factor BasR activates the *ors* gene cluster and suppresses (−) the expression of the emericellamide (*eas*) gene cluster. The involvement of AdaB and GcnE of the Saga/Ada complex has been experimentally proven (4).

## Discussion

### *S. rapamycinicus* induces a unique chromatin landscape in *A. nidulans*

By genome-wide ChIP-seq analysis of acetylated histone H3 (H3K9ac, H3K14ac) and the quantification of H3 we were able to uncover the chromatin landscape in the fungus *A. nidulans* upon co-cultivation with *S. rapamycinicus*. In an attempt to characterize the general distribution of nucleosomes and acetylation marks over the genome we compared the intensity of chromatin states with gene density. A lower gene density was typically found in heterochromatic regions such as the centromeres and telomeres creating a repressing environment (13). We found reduced H3 occupancy in heterochromatic regions indicating either replacement of H3 by the centromere-specific H3, CENP-A or reduced nucleosome occupancy (13, 14).

We observed distinct peaks for H3K9ac in *A. nidulans* grown in co-culture with *S. rapamycinicus*. One of the areas with the highest increase in H3K9ac was the *ors* gene cluster, nicely confirming our previous findings (4) (Fig. 7). Furthermore, previous ChIP qRT-PCR experiments indicated a distinct increase of H3K9ac inside the cluster borders, which did not expand to neighboring genes (4). By contrast, the H3K14ac modification seemed to be of a more global nature and not exclusively confined to specific regions such as the *ors* gene cluster. These conclusions were extended here by the pattern detected in the genome-wide ChIP-seq data, showing no spreading of H3K9ac to genes adjacent to the *ors* gene cluster which also demonstrates the quality of the genome-wide ChIP data generated here. Furthermore, these results are also consistent with our previous finding of reduced expression of SM cluster genes as a consequence of the lack of H3K14 acetylation (5). In contrast to H3K14ac, H3K9ac is less uniformly distributed over the genome. It only showed strong enrichment at promoters of certain genes. Especially high levels of acetylation were found at *orsA* and the bidirectional promoters of *orsD* and *orsE*. This observation was recently confirmed by the finding that H3ac and H3K4me3 were increased at the *orsD* gene only when the *ors* cluster was transcriptionally active (15).

We also assessed the distribution of H3K9ac and H3K14ac as well as the C-terminus of H3 (H3Cterm) at the TSS and translation termination sites (TTSs) (SI Results - Chromatin profiles at translation start sites and translation termination sites). For H3K9, an enrichment of acetylation ~500 bp downstream of the TSS as well as immediately upstream of the TSS was observed. This was expected as similar results were obtained with an antibody targeting the acetylated N-terminus of histone H3 in *A. nidulans* (15) and other fungi such as *S. cerevisiae* and *Cryptococcus neoformans* (16, 17). Increased acetylation coincides with reduced levels of H3 around the TSS, which is most likely due to a depletion of nucleosomes at the promoter. The profile plots for H3K14 acetylation are similar, although not as highly enriched around the TSS as H3K9 (Fig. S11 a). As expected, a comparison of LFCs for both modifications showed high similarity suggesting that they are established interdependently (18, 19). At the 3’ end of the ORF, H3 density drastically increased accompanied by reduced levels of H3K9ac and H3K14ac (Fig. S11). Likewise, reduced acetylation at the TTS was observed in *A. nidulans* (15) and *S. cerevisiae* (17). It is interesting to notice, that the increase in nucleosome density directly correlated with a decrease in the gene expression rate (Fig. S12 e). Previous studies suggested a direct correlation between the presence of nucleosomes and the stalling of RNA polymerase II (20).

### Increased gene expression directly correlates with histone H3K9 acetylation

Acetylation is generally regarded as an activating chromatin mark promoting transcription of eukaryotic genes (21). Our study suggests a more differentiated picture. When we compared data from this study with microarray data (4) (Figs. S3 & S4) the acetylation of H3K9 directly correlated with gene expression levels. A similar finding was reported for other fungi (22). By contrast, this was not observed for acetylation of H3K14. This could partly result from the low number of targets for this modification. By contrast, gene promoters showed a distinct increase of H3K14ac at the TSS in dependence on the average transcription level (Fig. S12 c). The low correlation between active gene transcription and acetylation at H3K14 confirmed earlier results. Previously, we showed that a mimicry of a hypo-acetylated lysine 14 on histone H3 drastically altered the phenotype and the expression of SM gene clusters in this strain (5). This effect, however, was overcome when later time points of cultivation were considered. Taken together, the primary location at the TSS and the major defect in SM production at earlier stages indicate a role for H3K14ac in transcriptional initiation. Hyper-acetylation at H3K14 could be also relevant for marking active genes and providing a docking side for regulatory proteins.

### *S. rapamycinicus* silences fungal nitrogen metabolism

A substantial number of genes involved in primary and secondary nitrogen metabolism were strongly depleted for H3K9ac upon co-cultivation. This correlated with reduced expression of the respective genes. Thus, upon contact with the bacterium, *A. nidulans* showed reduced nitrogen uptake and reduced degradation of various nitrogen sources, leading to nitrogen starvation.

Under nitrogen starvation or low availability of primary nitrogen sources, such as glutamine and ammonium, the intracellular level of glutamine drops (23). This was in fact observed for the intracellular concentration of amino acids in *A. nidulans* when the fungus was co-cultured with the bacterium (Fig. S13). Thus, in presence of *S. rapamycinicus* but not of non-inducing streptomycetes like *S. lividans* the fungus is in a physiological state of nitrogen starvation (Fig. 7). Nitrogen limitation was shown before to represent a trigger for the activation of a number of SM gene clusters including the *ors* gene cluster (24, 25). Nitrogen starvation also activates the expression of the anthrone *(mdp)* gene cluster (24), which we also observed in our data. However, induction of orsellinic acid production by nitrogen starvation took about 60 h, whereas co-cultivation with *S. rapamycinicus* already triggered expression of the cluster genes after 3 h. Therefore, it is unlikely that the bacteria-triggered activation of the cluster is exclusively achieved by restricting nitrogen availability for the fungus. Furthermore, shortage of nitrogen leads to de-repression of genes involved in the usage of secondary nitrogen sources, which was not supported by our data. In *S. cerevisiae*, it was shown that a shift from growth under nutrient sufficiency to nitrogen starvation induced degradation of mitochondria (26). Similarly, upon contact of *A. nidulans* with the bacterium decreased acetylation and transcription of genes with mitochondrial function were also detected. This was further supported by a lower mitochondrial respiratory activity in the fungal cells during co-cultivation (Fig. 3 c).

### BasR is a central regulatory node for integrating bacterial signals leading to regulation of SM gene clusters

Another consequence of nitrogen starvation is the reduced availability of amino acids in the cell. Consequently, as shown here, the amino acid biosynthetic pathways represented a major group of de-regulated genes at both the acetylation level and expression level. Amino acid biosyntheses in fungi are regulated by the CPC system upon starvation for distinct amino acids (23, 27). Since deletion of *cpcA* in *A. nidulans* did not affect the induction of the *ors* gene cluster, but on the other hand the artificial inducer of the CPC system 3-AT (28), it was conceivable that CPC somehow plays a role. 3-AT is a structural analogue of histidine triggering histidine starvation in the fungal cell and thereby the CPC (28). In *S. cerevisiae*, other regulators were also shown to induce the CPC such as the heterodimeric transcription factor complex Bas1/Bas2p (10, 29) which is even bound by Gcn5. We identified two putative orthologous genes in the genome of *A. nidulans*. Further analysis revealed only *basR (AN7174)* as being involved in the *ors* gene cluster activation during the fungal-bacterial co-cultivation (Fig. 7). Despite the fact that *AN8377* seems to be more similar to the *S. cerevisiae bas1* (Figs. 6 & S8), it is not needed for the *ors* gene cluster activation.

Based on bioinformatic analysis BasR of *A. nidulans* consists of 305 amino acids and thus is rather different from its closest homolog Bas1p of *S. cerevisiae* with 811 amino acids (30). The BIRD region of Bas1p mediating the Bas1p-Bas2p interaction (11), is missing in BasR. The *basR* gene was highly up-regulated in the microarray data which coincided with increased H3K9 acetylation of its promoter. *basR* deletion and overexpression clearly demonstrated the function of this transcription factor gene in activating the *ors* gene cluster in dependence of *S. rapamycinicus*. For efficient *basR* expression a functional GcnE seems to be required, indicating a similar dependency as observed for *bas1* in yeast (10).

Interestingly, the *basR* gene could not be found in all fungal genomes analyzed here, but for example in *A. sydowii* and *A. versicolor* which were also found to encode *ors* gene clusters. Like in *A. nidulans*, overexpression of the *A. sydowii basR* gene led to the activation of its silent *ors* gene cluster. Based on this finding we predicted that *S. rapamycinicus* also induces the *ors* gene cluster in *A. sydowii* which indeed was the case. We did not find a *basR* homologue in *A. fumigatus* although the formation of fumicyclines is induced by *S. rapamycinicus* (31). This might be due to the genome data available that lack the *basR* gene due to missing annotation or, alternatively, a different regulatory response mechanism to *S. rapamycinicus* is present in *A. fumigatus*.

Genome-wide ChIP-seq analysis also indicated that the interaction of *S. rapamycinicus* with *A. nidulans* influenced other SM gene clusters, *e.g*., it led to repression of the formation of emericellamides. Also for this repression, BasR was required, indicating that overexpression of *basR* phenocopies the regulation by *S. rapamycinicus*. As implied by the finding that the presence of *basR* and the *ors* cluster in several fungi coincided with their inducibility by *S. rapamycinicus*, in future it might be possible to predict which microorganisms talk to each other based on their genetic inventory.

## Material and Methods

### Microorganisms, media and cultivation

Microorganisms are listed in Table S4. *A. nidulans* strains were cultivated in *Aspergillus* minimal medium (AMM) at 37 °C, 200 rpm (32). When required, supplements were added as follows: arginine (871 μg/mL), p-aminobenzoic acid (3 μg/mL) and pyridoxine HCl (5 μg/mL). Pre-cultures were inoculated with 4 × 10^8^ spores per mL. 10 μg/mL doxycycline was used to induce the *tet*^On^ inducible system. For the measurement of orsellinic acid, mycelia of overnight cultures (~16 h) in AMM were transferred to fresh medium and inoculated with *S. rapamycinicus*, as previously described (3). RNA extraction for expression analysis during co-cultivation was performed after 3 hours of cultivation, for analysis of the *basR* overexpression mutant after 6 hours of monoculture; samples for HPLC analysis were taken after 24 h. *A. sydowii* was cultivated at 28 °C, 200 rpm in malt medium (33). For the induction of the *ors* cluster in *A. sydowii*, 48-hour old precultures were transferred to fresh AMM and inoculated with *S. rapamycinicus* or doxycycline. 10 μg/mL doxycycline was added twice over the course of 48 hours. Samples were taken for LC-MS analysis after 96 hours for *A. sydowii* co-cultivation and after 48 hours for the *A. sydowii basR* overexpression mutant. For MALDI-MS Imaging analysis conductive ITO slides (Bruker Daltonics, Bremen, Germany) were coated with 3 mL 0.5% (w/v) AMM agar and incubated at room temperature for 30 minutes (34, 35). Identical conditions were ensured by supplementation of all slides with arginine regardless of the fungal genotype. *S. rapamycinicus* was applied by filling 5 mL of a preculture in a tube and point inoculation of 15 μl of the settled mycelium on the agar. For *A. nidulans*, 500 conidia of wild type and mutants were point inoculated on the agar. For co-cultivation experiments, both microorganisms were inoculated 1 cm apart from each other. The slides were incubated at 37 °C in a petri dish for 4 days. The slides were dried by incubation in a hybridization oven at 37 °C for 48 hours.

### Quantitative RT-PCR (qRT-PCR)

Total RNA was purified with the Universal RNA Purification Kit (roboklon, Berlin, Germany). Reverse transcription of 5 μg RNA was performed with RevertAid Reverse Transcriptase (Thermo Fisher Scientific, Darmstadt, Germany) for 3 hours at 46 °C. qRT-PCR was performed as described before (3). The *A. nidulans* ß-actin gene (*AN6542*) served as an internal standard for calculation of expression levels as previously described (3). Primers for amplification of probes are listed in Table S5.

### Preparation of chromosomal DNA and Southern blot analysis

*A. nidulans* genomic DNA was isolated as previously described (3). Southern blotting was performed using a digoxigenin-11-dUTP-labeled (Jena Bioscience, Jena, Germany) probe (3).

### ChIP coupled to qRT-PCR analysis

ChIP coupled to qRT-PCR analysis including crosslinking of the DNA, sonication, antibody incubation and precipitation and DNA reverse-cross-linking is described in SI Material and Methods.

### Extraction of fungal compounds, HPLC and LC-MS analyses

Extraction of *A. nidulans* and *A. sydowii* monocultures as well as co-cultures with *S. rapamycinicus* and the subsequent HPLC and LC-MS analysis are described in SI Material and Methods.

### MALDI-MS imaging analysis and data processing

Sample preparation and matrix coating were performed as previously described (34). Samples were analyzed (34) in an UltrafleXtreme MALDI TOF/TOF (Bruker Daltonics, Bremen, Germany), in reflector positive mode with the following modifications: 100-3000 Da range, 30 % laser intensity (laser type 4) and raster width 200 μm. The experiments were repeated three times (2^nd^ and 3^rd^ replicates with 250 μm raster width). Calibration of the acquisition method, spectra procession, visualization, analysis and illustration were performed as described before (34). Chemical images were obtained using Median normalization and weak denoising.

### Resazurin assay

Respiratory activity was measured by reduction of resazurin to the fluorescent dye resorufin. 10^4^ conidia of *A. nidulans* in 100 μL AMM were pipetted in each well of a black 96 well plate. The plate was incubated for 16 hours at 37 °C. The pre-grown fungal mycelium was further cultivated in monoculture or with 10 μL of an *S. rapamycinicus* culture. Cultures were further supplemented with 100 μL of AMM containing resazurin in a final concentration of 0.02 mg/mL. Fluorescence was measured (absorption wavelength 560 nm, emission wavelength 590 nm) every 30 minutes for 24 hours at 37 °C in a Tecan Fluorometer (Infinite M200 PRO, Männedorf, Switzerland). For all conditions, measurements were carried out in triplicates for each of the two biological replicates. Significance of values was calculated using 2-way ANOVA Test with GraphPad Prism 5 (GraphPad Software Inc., La Jolla, USA).

**ChIP-seq pre-processing, DCS analysis and MACS analysis** are described in SI Material and Methods.

**Generation of *A. nidulans* and *A. sydwoii* mutant strains** is described in SI Material and Methods.

### Phylogenetic analysis

The amino acid sequences for the two Myb-like transcription factors from *A. nidulans* (*AN7174* (*basR*) and *AN8377*) and Bas1 from *S. cerevisiae* were used for a Blast search in the UniProtKB database. For each sequence, the first 50 hits were retrieved. All hits were grouped together, and redundant and partial sequences removed. The obtained 54 hits were firstly aligned using MUSCLE (36). The phylogenetic tree was obtained using the Maximum Likelihood method contained in the MEGA6 software facilities (37).

## Availability of data and materials

ChIP-seq data were deposited in the ArrayExpress database at EMBL-EBI (www.ebi.ac.uk/arrayexpress) under accession number E-MTAB-5819. The code for data processing and analysis can be obtained from https://github.com/seb-mueller/ChIP-Seq_Anidulans.

## Acknowledgements

We thank Christina Täumer and Karin Burmeister for excellent technical assistance and Sven Krappmann (Friedrich-Alexander University, Erlangen-Nürnberg, Germany) for kindly providing plasmid pSK562. Financial support by the Deutsche Forschungsgemeinschaft (DFG)-funded excellence graduate school Jena School for Microbial Communication (JSMC), the International Leibniz Research School for Microbial and Biomolecular Interactions (ILRS) as part of the JSMC, the DFG-funded Collaborative Research Center 1127 ChemBioSys (projects B01, B02 and INF), the BMBF-funded project DrugBioTune in the frame of Infectcontrol2020 and the European Research Council for a Marie Skłodowska-Curie Individual Fellowship (IF-EF; Project reference 700036) to María García-Altares is gratefully acknowledged.

## Supplementary Information

### SI Material and Methods

#### ChIP coupled to quantitative RT-PCR (qRT-PCR)

Cultures were grown as described in the cultivation part. After 3 hours the isolated DNA was cross-linked to proteins as described before (1). Powdered mycelium was dissolved in 1 mL of sonication buffer (1) and 330 μL aliquots were then subjected to sonication for 30 min with cycles of 2 min maximum intensity followed by a 1 min pause. Sheared chromatin was separated from cell wall debris and incubated with 40 μL of a protein A slurry for 30 min at 4 °C on a rotary shaker. A purified 1:10 dilution of the supernatant was then incubated overnight at 4 °C with 3 μL of antibody directed against the desired target. Antibodies were precipitated with 40 μL of Dynabeads (Invitrogen, Carlsbad, USA) and were immediately incubated with the sample for 40 min at 4 °C on a rotary shaker. Samples were washed 3 times with low salt buffer (1) followed by one time washing with high salt buffer (1). Washed beads were dissolved in 125 μl TES buffer and reverse cross-linked with 2 μL of 0.5 M EDTA, 4 μL of 1 M Tris-HCl pH 6.5 and 2 μL of 1 mg/mL proteinase K for 1 hour at 45 °C. Subsequent DNA purification was conducted with a PCR purification kit and samples were eluted in 100 μL of 1:10 diluted elution buffer. The DNA concentration of genes of interest was quantified using qRT-PCR as described above. Antibodies used are the following: mouse monoclonal ANTIFLAG M2 (Sigma-Aldrich, F3165-5MG, Taufkirchen, Germany), rabbit polyclonal anti-histone H3 (Abcam 1791, Cambridge, UK), rabbit polyclonal histone H3K9ac (Active Motif, Catalog No: 39161, La Hulpe, Belgium) and rabbit polyclonal anti-acetyl-histone H3 (Lys14) (Merck Millipore, Darmstadt, Germany).

#### ChIP-seq pre-processing

The *A. nidulans* FGSC A4 genome and annotation (version s10-m03-r28) were obtained from the *Aspergillus* Genome Database (AspGD) (2). The *S. rapamycinicus* NRRL 5491 genome was obtained from NCBI (GI 521353217). Both genomes were concatenated to a fused genome which served as the reference genome for subsequent mapping. Raw ChIP-seq reads were obtained by using FastQC v0.11.4. Trimming and filtering were achieved by applying Trim Galore utilizing Illumina universal adapter and phred+33 encoding. Reads were not de-duplicated since the duplication rate was < 15% for most libraries. Bowtie2 (version 2.2.4) using default parameters was employed to map reads to the fused genome. Quantification of reads was carried out using the Bioconductor ‘GenomicAlignments’ package forming the basis for three subsequent approaches. Firstly, a genome-wide equi-spaced binning across the genome with different resolutions (50k and 2k bp bins) counting reads overlapping each bin was applied. Library normalization on bin counts was performed by only considering reads mapping to the *A. nidulans* genome. Secondly, reads overlapping genes were counted, using the AspGD (2) annotation. They formed the basis for the subsequent DCS analysis (see below). Thirdly, average profile plots to assess relative histone distributions around TSS and TTS were generated using the bioconductor package regioneR(3).

#### DCS analysis

To identify genes exhibiting differences in their chromatin state, we employed the bioconductor package edgeR (4) originally developed for RNA-seq differential expression analysis. The ChIP-seq data follow the same pattern, *i.e*., negative binomial distribution of reads. Library normalization was achieved with the trimmed mean of M values (4) method only based on *A. nidulans* gene counts for calculating the effective library sizes, not taking into account reads mapping to *S. rapamycinicus* which would otherwise artificially influence the effective library size. Comparisons were made between libraries for all ChIP targets separately obtained from monocultures of *A. nidulans* and co-cultures with *S. rapamycinicus*. These targets were H3, H3K9ac and H3K14ac. Results including normalized read counts (RPKM) statistics and LFCs are reported in Table S2. Normalized counts and LFCs were also further used for comparisons with the corresponding microarray-based gene expression and the calculated LFCs.

#### MACS analysis

Candidate peaks were identified using two methods: a differential binding analysis (EdgeR) and a peak-calling approach (MACS, version 2.0.1) (5). The peak caller performed several pairwise comparisons between samples with the same antibody and different conditions in order to retrieve the peaks with significant change of ChIP signal indicating differential binding for that particular comparison. The program kept the track of different replicates, the signal was reported per million reads and produced a BED format track of the enriched regions, other parameters were used with default values. The BED files were subsequently converted to Big Wig format for visualization through the tool Integrative Genomics Viewer(6).

#### Extraction of fungal compounds, HPLC and LC-MS analyses

Culture broth containing fungal mycelium with and without bacteria was homogenized utilizing an ULTRA-TURRAX (IKA-Werke, Staufen, Germany). Homogenized cultures were extracted twice with 100 mL ethyl acetate, dried with sodium sulfate and concentrated under reduced pressure. For HPLC analysis, the dried extracts were dissolved in 1-1.5 mL of methanol. Analytical HPLC was performed using a Shimadzu LC-10Avp series HPLC system composed of an autosampler, high pressure pumps, column oven and PDA. HPLC conditions: C18 column (Eurospher 100-5 250 × 4.6 mm) and gradient elution (MeCN/0.1 % (v/v) TFA (H_2_O) 0.5/99.5 in 30 min to MeCN/0.1 % (v/v) TFA 100/0, MeCN 100 % (v/v) for 10 min), flow rate 1 mL min^−1^; injection volume: 50 μL.

The samples of *A. sydowii* were loaded onto an ultrahigh-performance liquid chromatography (LC)-MS system consisting of an UltiMate 3000 binary rapid-separation liquid chromatograph with photodiode array detector (Thermo Fisher Scientific, Dreieich, Germany) and an LTQ XL linear ion trap mass spectrometer (Thermo Fisher Scientific, Dreieich, Germany) equipped with an electrospray ion source. The extracts (injection volume, 10 μL) were analyzed on a 150-by 4.6-mm Accucore reversed-phase (RP)-MS column with a particle size of 2.6 μm (Thermo Fisher Scientific, Dreieich, Germany) at a flow rate of 1 mL/min, with the following gradient over 21 minutes: initial 0.1% (v/v) HCOOH-MeCN/0.1% (v/v) HCOOH-H_2_O 0/100, which was increased to 80/20 in 15 min and then to 100/0 in 2 min, held at 100/0 for 2 min, and reversed to 0/100 in 2 minutes.

Identification of metabolites was achieved by comparison with an authentic reference. Samples were quantified *via* integration of the peak area using Shimadzu Class-VP software (version 6.14 SP1).

#### Generation of *A. nidulans* deletion strains

The transformation cassettes for the *basR* and *AN8377* deletion strains were constructed as previously described (7). Approximately ~1000-bp sequences homologous to the regions upstream and downstream of *basR* and *AN8377* were amplified and fused to the *argB* deletion cassette (8). Transformation of *A. nidulans* was carried out as described before (9).

#### Generation of inducible *A. nidulans* and *A. sydowii basR* overexpressing mutant strains

For overexpression of *basR*, the tetracycline-controlled transcriptional activation system (*tet*^On^) was used (10). The *basR* gene sequences together with their ~1000-bp flanking regions were amplified from *A. nidulans* and *A. sydowii* genomic DNA. The *tet*^On^-system was amplified from plasmid pSK562. All DNA fragments were assembled by using NEBuilder HiFi DNA Assembly Master Mix (New England Biolabs, Frankfurt, Germany). The *A. nidulans pabaA1* gene was used as a selectable marker to complement the p-aminobenzoic acid auxotrophy of the *A. nidulans* Δ*basR* mutant. For *A. sydowii*, the *Aspergillus oryzae hph* cassette was used as the selectable marker. 200 μg/mL hygromycin (Invivogen, Toulouse, France) were used for selection of transformant strains.

#### Measurement of amino acids

Amino acids were extracted from 10 mg samples with 1 mL of methanol and the resulting extract was diluted in a ratio of 1:10 (v:v) in water containing the ^13^C, ^15^N labeled amino acid mix (Isotec, Miamisburg, Ohio, USA). Amino acids in the diluted extracts were directly analyzed by LC-MS/MS as described with the modification that an API5000 mass spectrometer (Applied Biosystems, Foster City, California, USA) was used (11).

### SI Results

#### Details of the ChIP analysis

After first examination, we found that a significant proportion of co-incubated library reads originated from *S. rapamycinicus*. A fused genome concatenating the *A. nidulans* and the *S. rapamycinicus* genomes was generated. About 90-98% of the reads mapped against the fused genome (Table S2), which suggested a high quality of sequencing data. This assumption was confirmed by examining the quality of libraries using FastQC (data not shown). As indicated, the coverage was substantially higher on the *S. rapamycinicus* genome with only ~20 % mapping to the *A. nidulans* genome (see Table S2). Expectedly, this ratio has shifted considerably towards the *A. nidulans* genome for histone targeting ChIP libraries (Fig. 1a) going up from 20 % to around 90 % for all H3, H3K9 and H314 libraries validating correct antibody enrichment as *S. rapamycinicus* is devoid of histones. However around 10 % of reads were still mapping to the *S. rapamycinicus* genome which might be due to imperfect antibody specificity. Coverage depth deviations were accounted for by only considering reads originating from *A. nidulans*. This allowed for calculation of library size factors used for library normalization. To assess antibody specificity, we calculated the fraction of reads mapping to mitochondria, which do not contain histones. The control library amounted to about 0.25 % of reads as opposed to about 0.01-0.03 % for H3, H3K9ac, H3K14ac libraries constituting a 10-fold enrichment. As a background control we used ChIP material obtained from anti-FLAG-tag antibody precipitates of a non-tagged fungal wild-type strain co-cultivated under the same conditions with *S. rapamycinicus*.

We quantified the relative library proportions which amounted to 8-17 % of H3, H3K14ac and H3K9ac as well as up to ~65 % for background reads of co-incubated libraries mapped to *S. rapamycinicus* (Table S2). However, the background read proportions might not necessarily reflect actual gDNA ratios of both species in the co-cultivation due to various potential biases. To examine read distribution for each library, we counted mapped reads within equally spaced bins along the fused genome for different resolutions (see methods and Fig. 1 a & b). As expected, background reads (upper panel of Fig. 1 a) were evenly distributed across the genome reflecting nonspecific targeting of particular areas. The fused genome further enabled for controlling correct co-incubation conditions since no reads should be mapping to *S. rapamycinicus* in non-co-incubated samples as can be seen in Fig. 1 a in the right panels (blue lines). The co-incubated samples exhibit an increased coverage in the middle of the *S. rapamycinicus* genome which might be due to sequence biases or enriched DNA caused by the replication origin (*oriC*) located in this region (12). Further, there were coverage dips in the middle of the fungal chromosomes (see Fig. 1 a), which were most likely due to incomplete assembly around the centromeres which are characterized by long ‘N’ stretches (13).

#### Changes of H3K9 and H3K14 acetylation profiles in *A. nidulans* in response to *S. rapamycinicus*

As reported in the manuscript, the genome-wide H3K9 and H3K14 chromatin landscape of *A. nidulans* was determined. There was also a drop-off in all libraries at the chromosome arms, which was most likely caused by the bordering bins being shorter and therefore account for less reads. Since gene density also varies across the genome (lower panel of Fig. 1 a), we addressed the question whether this correlates with the intensity of the investigated chromatin states. To this end, we calculated the spearman correlation to correlate the read counts and the gene counts among the 50k bins. As expected, there was almost no correlation between the background and the genes (r = 0.09). However, for H3 we found it to be rather high (r = 0.37) (Fig. S4). Since the used bin size is large, this could point at global H3 occupancy to be higher for high gene density regions such as euchromatin and low H3 occupancy for heterochromatin. Noteworthy, the correlation between read and gene density was found to be lower for H3K14ac and H3K9ac (r=0.14 and 0.15 respectively), which might indicate a more subtle regulatory mechanism for those marks targeting individual genes as opposed to larger domains. Notably, the highest correlation was found between the two acetylation marks (r=0.53) hinting at some potential cross-talk or common regulation between them (Fig. S5).

#### Chromatin profiles at translation start sites and translation termination sites

We assessed the location of H3K9ac, H3K14ac and histone H3 relative to promoters and gene bodies by plotting the average read count frequency for all genes to either the TSS or TTS (Fig. S1) (14). Due to missing information about the 5’ and 3’ transcriptional start and stop sites, we used translational start and stop sites for transcription start and stop sites, respectively, as surrogates. The results obtained apply to both *A. nidulans* in mono- and co-culture with the bacterium. According to Kaplan et al. (15), the peaks correspond to highly positioned nucleosomes relative to the TSS with a nucleosome-free region (NFR) directly upstream of the TSS. H3K9ac and H3K14ac showed highest enrichment for the first and second nucleosomes up- and downstream of the TSS and drastic reduction downstream of the third nucleosome after the TSS (Fig. S1). Reduced occupancy of unmodified histone H3 was observed at the TSS(15). Towards the 3’ end of genes, histone H3 occupancy gradually increased, which was accompanied by a decrease in acetylation. Plotting of differentially acetylated H3K9ac against H3K14ac showed a strong correlation between the localization of the two modifications (Fig. S14 a). Acetylation is generally described as an transcription activating mark(16). To test for this general assumption we correlated our acetylation data to microarray data generated under the same condition(17) (Fig. S14 b). This allowed us to compare the log-fold changes (LFCs) of the differential chromatin states with the LFCs calculated from the RNA expression data. H3K9ac correlates to the differentially expressed genes with a coefficient of 0.5 in contrast to H3K14ac (−0.05) and histone H3 (−0.01) for which no detectable dependency was determined (Fig. S3). Similarly, Fig. S3 shows the same trend, i.e., a correlation of 0.2 for gene expression changes *versus* H3K9ac changes which is expected since the calculation included all genes of which most did not change. To determine the correlation of acetylation at the TSS and the TTS according to the grade of expression of the genes, we separated the differentially expressed genes into four quartiles (q1 lower 25 %, q2 the medium lower 25 – 50 %, q3 are the medium higher 50-75 %, q4 higher 25 %). The increase of acetylation at the TSS correlated with the expression level of genes (Fig. S12 a & c). Decreased expression coincided with an increase of histone H3 at the TSS (Fig. S12 e).

#### Reduced intracellular amino acid concentration in response to the bacterium

The increased expression of *cpcA* and other genes involved in amino acid biosynthesis implied a reduced availability of amino acids in the cell upon bacterial-fungal co-cultivation. Therefore, we measured the internal amino acid pool in *A. nidulans* both grown in monoculture and with *S. rapamycinicus* (Fig. S13). As an additional control and to further confirm the specificity of the interaction, the fungus was co-cultivated with *Streptomyces lividans*, which does not induce the *ors* gene cluster. As shown in Fig. S13, significantly reduced levels of glutamine, histidine, phenylalanine, asparagine, threonine and reduced metabolism of arginine, which was supplemented to the medium, were observed. The monoculture of *A. nidulans*, co-cultivation of *A. nidulans* with *S. lividans* as well as the addition of *S. rapamycinicus* after 24 hours of fungal cultivation served as controls. All of the controls showed comparable amino acid levels.

**Figure S1.**
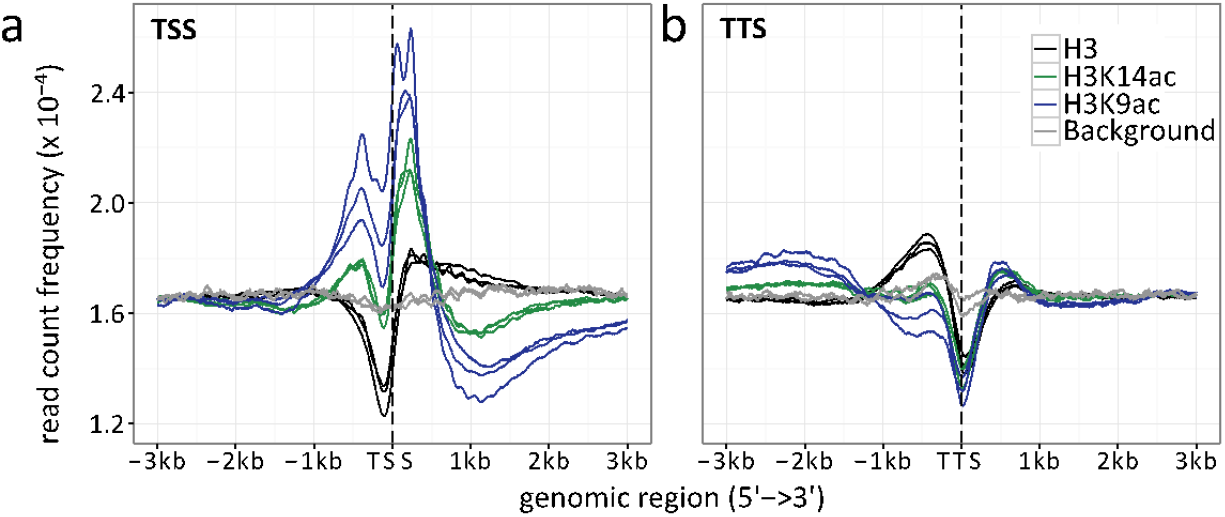
Read count frequencies for (a) TSS and (b) TTS. Lines correspond to the relative enrichment of ChIP signal strength relative to the TSS/TTS averaged across all genes. ChIP-seq read count serves as a surrogate for signal strength (see methods for further details). Compared were the enrichment of histone H3 (black), H3K9ac (blue), H3K14ac (green) and the background control (grey) over an average of all TSS and TTS. The enrichment curves for all biological replicates are given, indicated by multiple lines per enrichment target.

**Figure S2.**
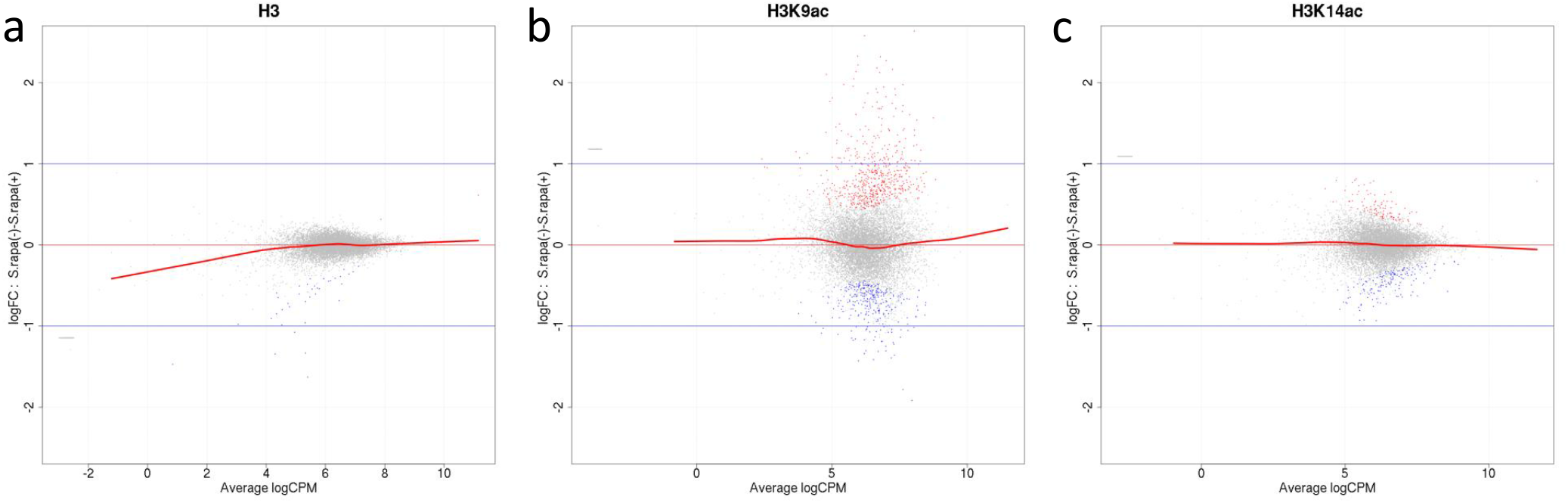
Mean average (MA) plots comparing normalized ChIP-seq LFCs H3(Cterm) (a), H3K14ac (b) and H3K9ac (c) of each gene between *A. nidulans* monoculture and co-culture for antibodies used in this study. Y-axis indicates LFCs of ChIP signal between mono- and co-culture for each gene which corresponds to the dots. X-axis indicates mean intensity of ChIP signal of both conditions. Genes were colored according to DCS outcome with grey indicating genes with no significant change of ChIP signal, red and blue indicate genes showing respective higher or lower signalin co-culture.

**Figure S3.**
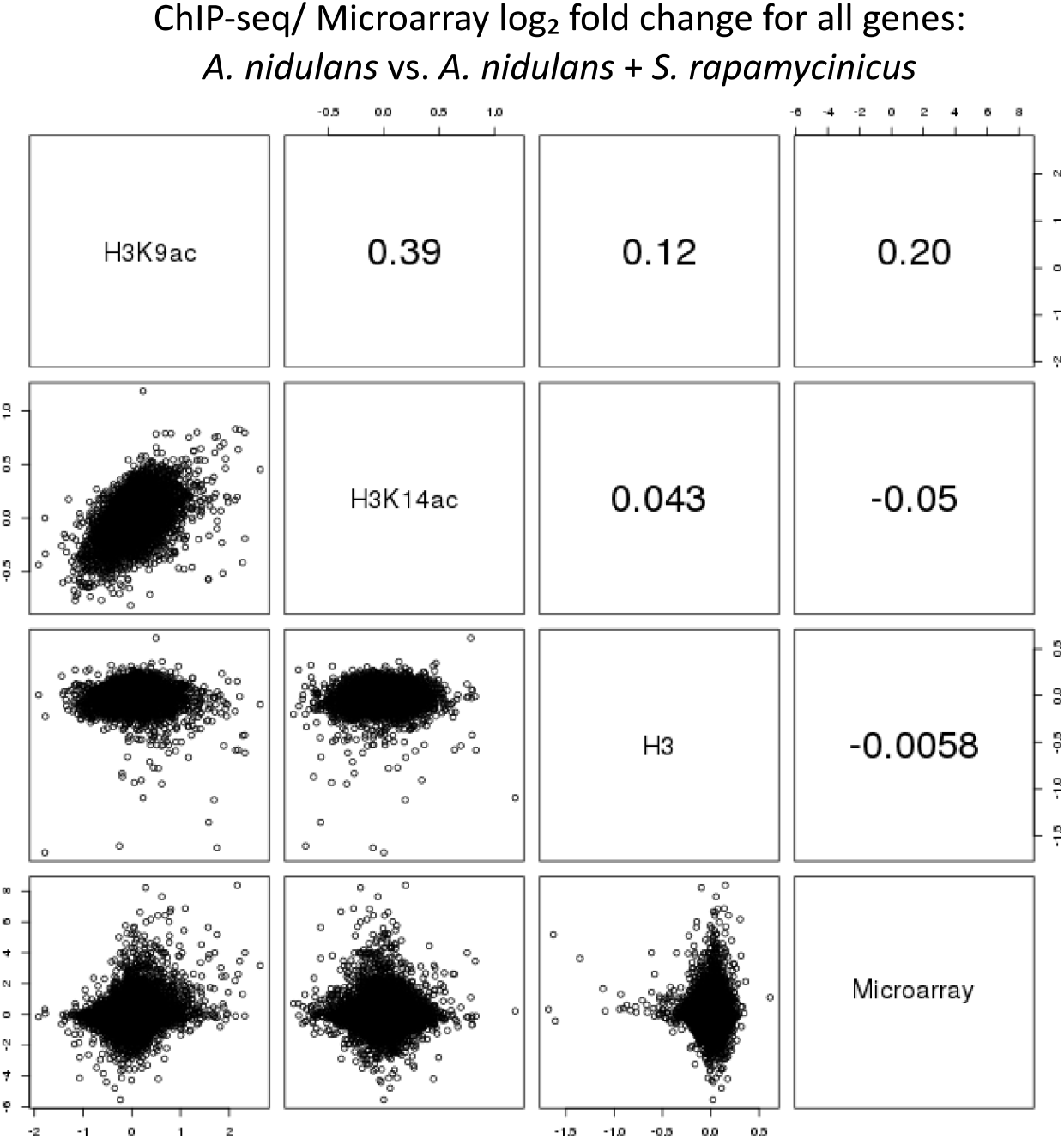
Correlation of data points for LFCs of ChIP-seq with LFCs of microarray data for all *A. nidulans* genes, depicting single data points and the correlation coefficient.

**Figure S4.**
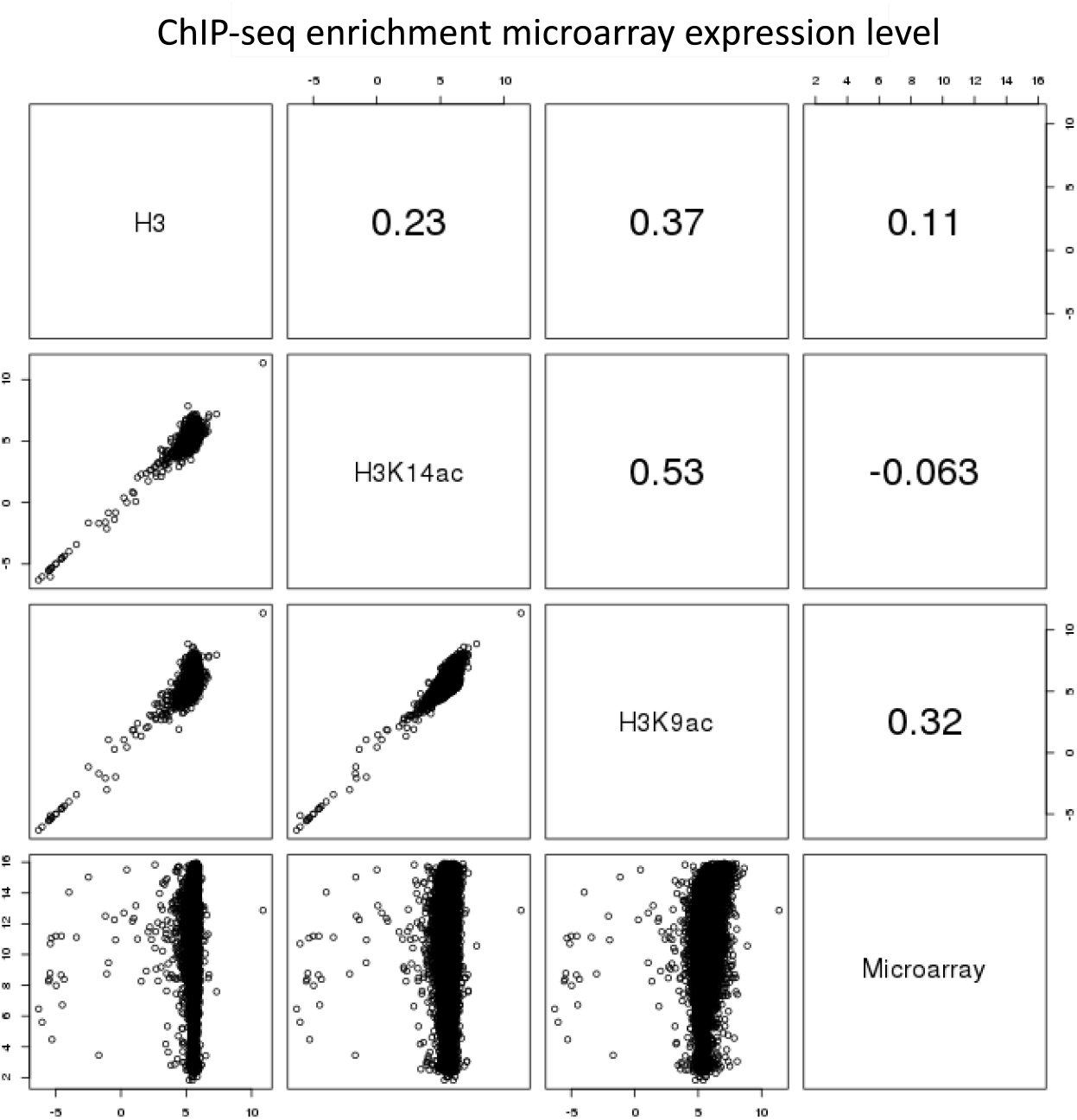
Pairwise comparison of ChIP-seq and microarray intensities of all genes in *A. nidulans* monoculture. The numbers resemble the correlation coefficient for the respective comparison. Intensity defines enrichment of number of reads per gene.

**Figure S5.**
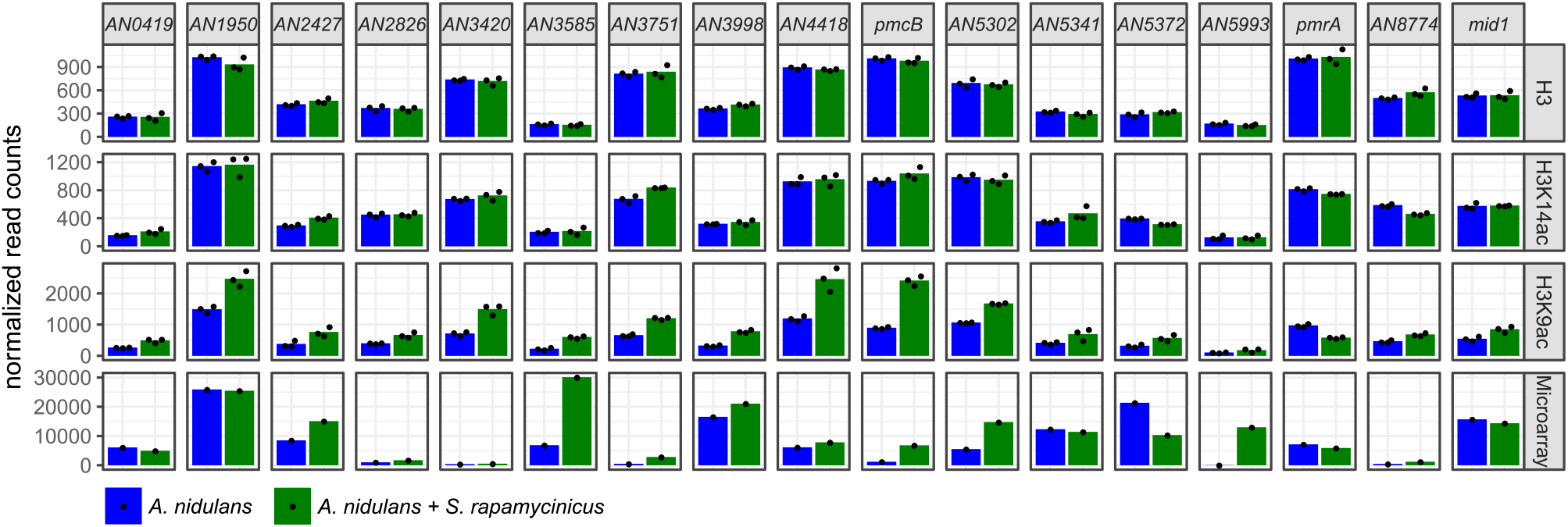
Normalized ChIP-seq read counts were used to quantify the chromatin state for individual genes. Here, for H3, H3K14ac and H3K9ac libraries genes involved in calcium signaling are shown. Counts were obtained by counting reads mapping to the promoter area for each gene that is 500 bp down- and 1000 bp upstream from the TSS. Depicted bars are calculated from three data points.

**Figure S6.**
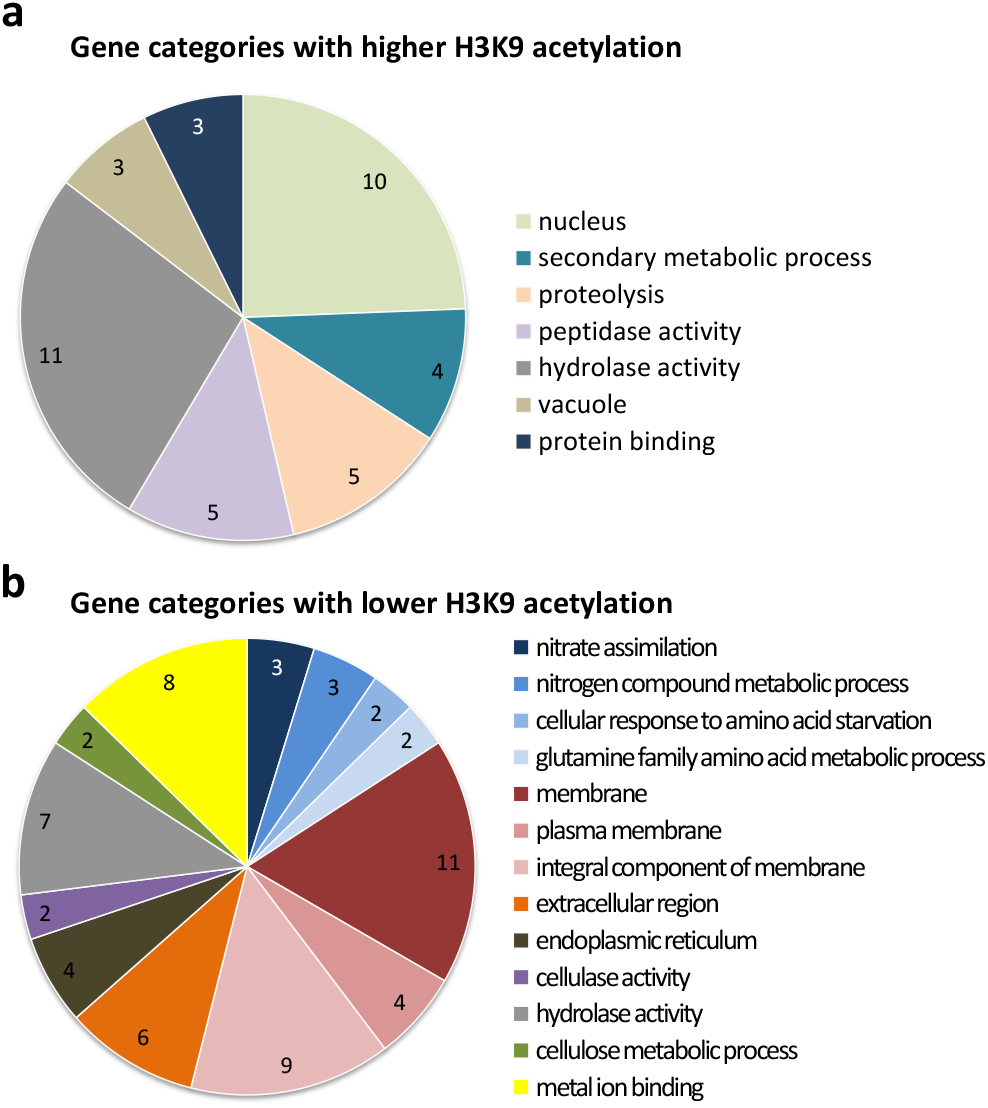
Gene ontology of the 15 most significantly enriched categories for differentially higher and lower acetylated genes at H3K9 upon co-cultivation with *S. rapamycinicus*. Functional categorization of differentially higher (A) and lower (B) acetylated genes, possessing a *p* value < 0.05, with FungiFun2 (18). Overrepresented categories having a p value < 0.01.

**Figure S7.**
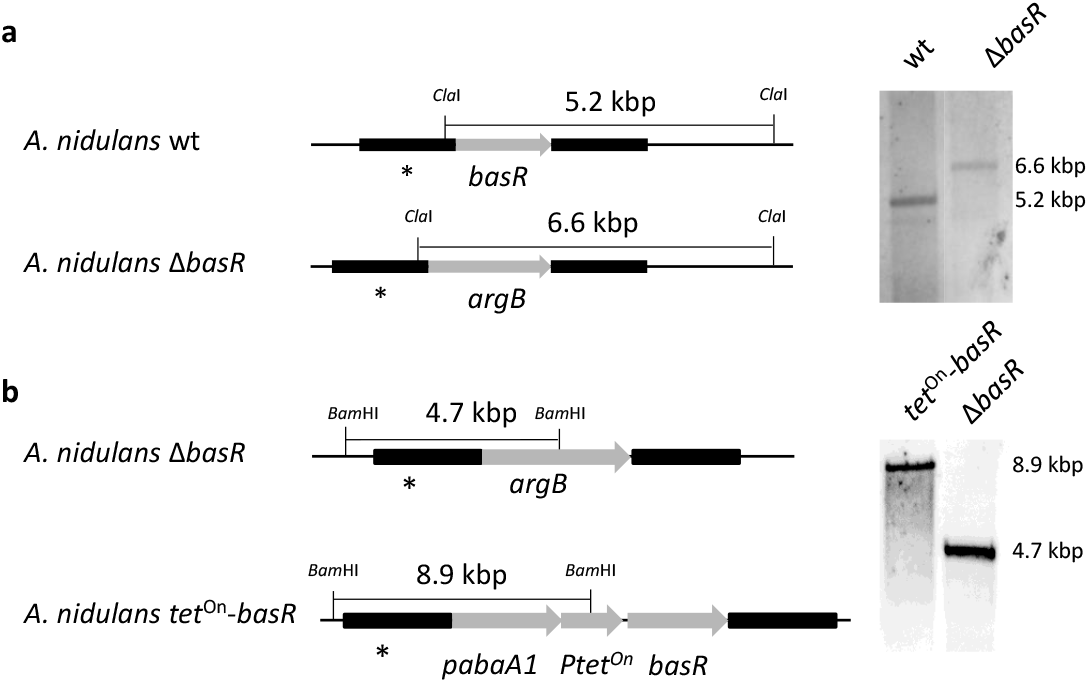
Generation of a *basR* deletion mutant and inducible overexpression strain based on the *A. nidulans* wild-type strain A1153. (**a**) Genomic organization of *basR* and Southern blot analysis of *basR* deletion. The *basR* gene was replaced by the *argB* gene. Transformant strains were checked with a probe (*) directed against the flanking region of the construct. Genomic DNA was digested with *ClaI*. wt, wild-type strain as a control. (**b**) Generation of the inducible *basR* overexpression strain by complementation of the *basR* deletion strain. The *tet*^On^-*basR* gene cassette was integrated at the Δ*basR* genomic locus using the *pabA1* gene as selectable marker replacing the *argB* marker. Genomic DNA was cut with BamHI. Transformant strains were checked with a probe (*) directed against the flanking region of the construct.

**Figure S8.**
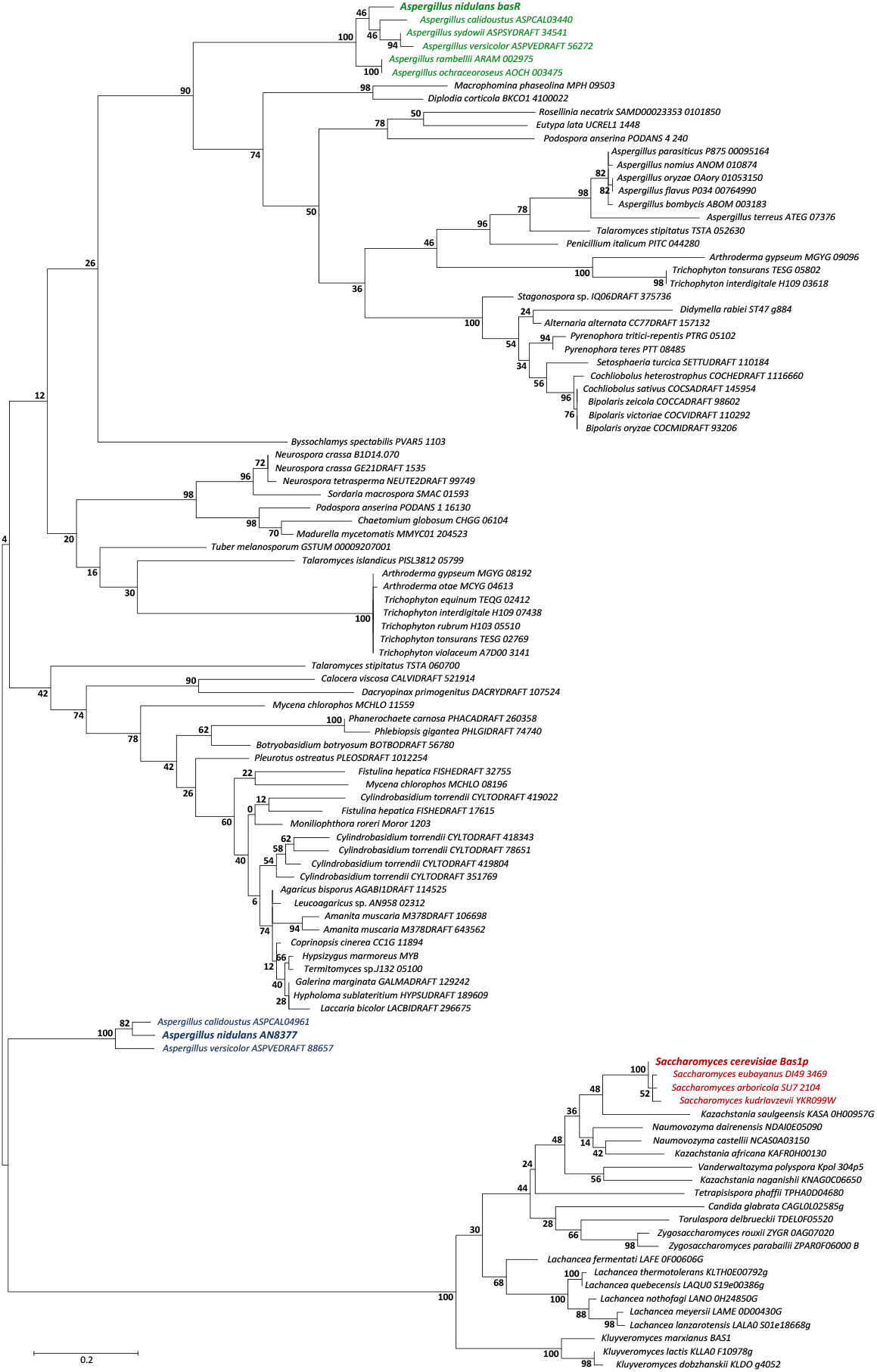
Molecular phylogenetic analysis of BasR (*AN7174*). The tree reports distances between BasR-similar amino acid sequences identified by BlastP analysis using the entire sequences. The percentage of trees in which the associated taxa clustered together is shown next to the branches. The BasR proteins from *A. nidulans, A. calidoustus, A. sydowii, A. versicolor, A. rambellii* and *A. ochraceoroseus* form a separate clade (reported in green), while the yeast Bas1p-related sequences are more distantly related to BasR (in red). The second similar Myb-like transcription factor from *A. nidulans* (*AN8377*) forms a clade with orthologues from *A. calidoustus* and *A. versicolor* (in blue), which seems to be more related to Bas1p than to BasR. The names of the selected sequences are given according to their UniProt accession numbers.

**Figure S9.**
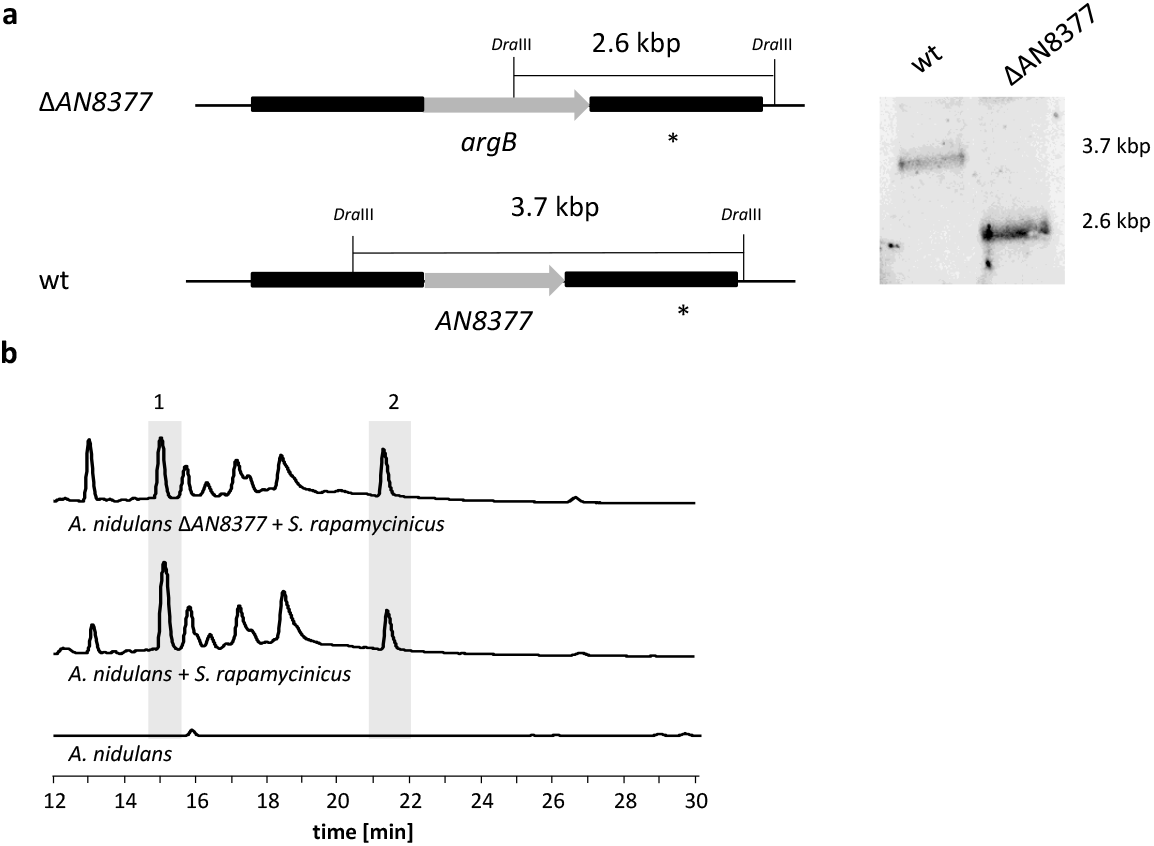
Deletion of the second putative *bas1p* homologous gene (*AN8377*) in *A. nidulans* and analysis of its impact on the *ors* gene cluster induction in response to *S. rapamycinicus*. (**a**) Chromosomal organization of the *A. nidulans AN8377* gene before and after deletion. The gene *AN8377* was replaced by an *argB* cassette in *A. nidulans* wild-type strain A1153. Genomic DNA was digested with DraIII. A PCR fragment covering the downstream sequence of *AN8377* was used as a probe (*). wt, wild-type strain as a control. (**b**) LC-MS-based detection of orsellinic acid (1) and lecanoric acid (2) in the co-cultivation of the *AN8377* deletion mutant with *S. rapamycinicus*.

**Figure S10.**
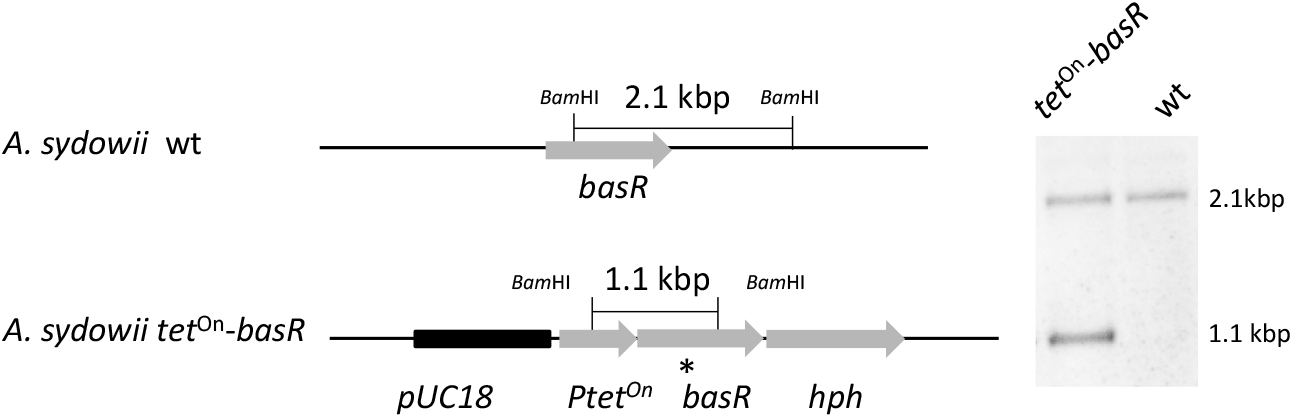
Generation of the inducible *basR* overexpression strain by ectopic integration of an additional copy of the *basR* gene in the *A. sydowii* wild-type strain (wt). The *tet*^On^-*basR* construct was integrated ectopically into the wild-type genome, using the *hph* cassette as selectable marker. For Southern blot analysis, transformant strains were checked with a probe (*) directed against a region flanking the *tet*^On^ cassette and *basR* gene. The genomic DNA was digested with BamHI.

**Figure S11.**
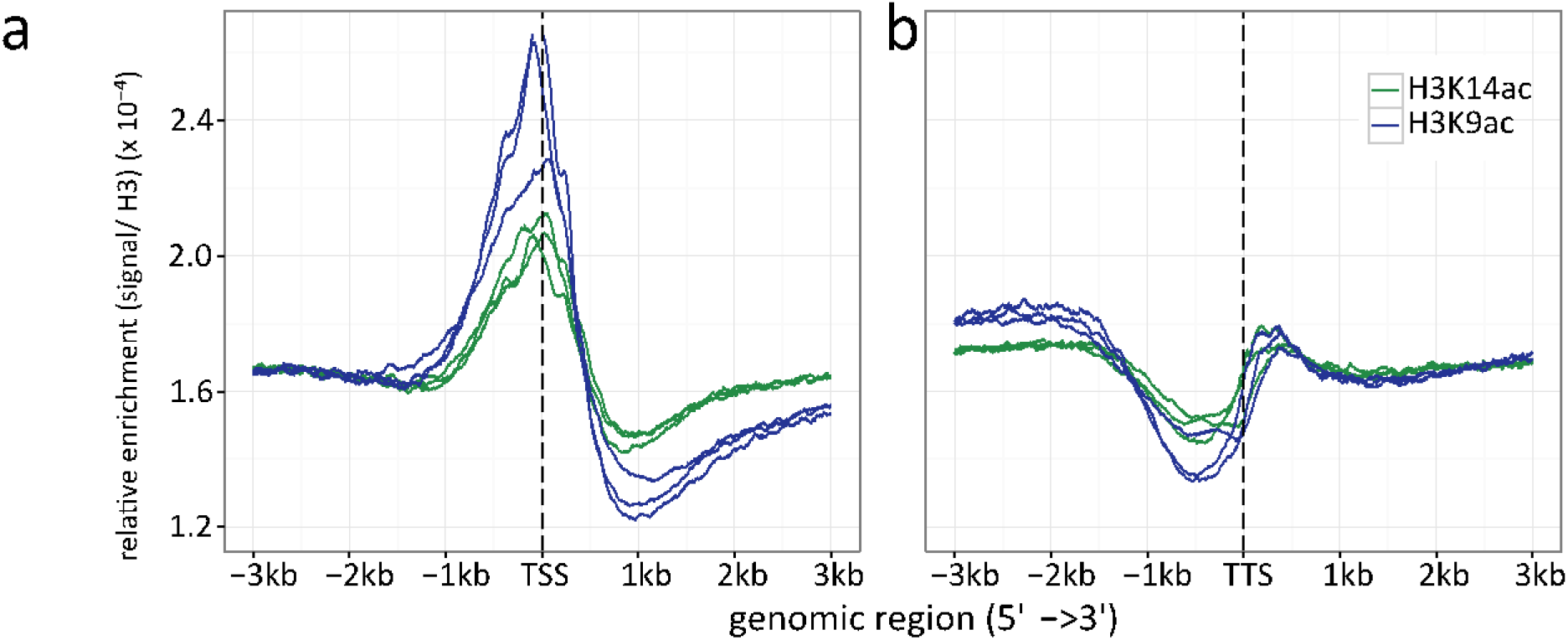
Histone H3 normalized read count frequencies for H3K9ac (green) and K14 ac (blue) at the (a) TSS and (b) TTS. The enrichment is given in signal to H3 ratio. Multiple lines per ChIP target resemble the three independent biological replicates.

**Figure S12.**
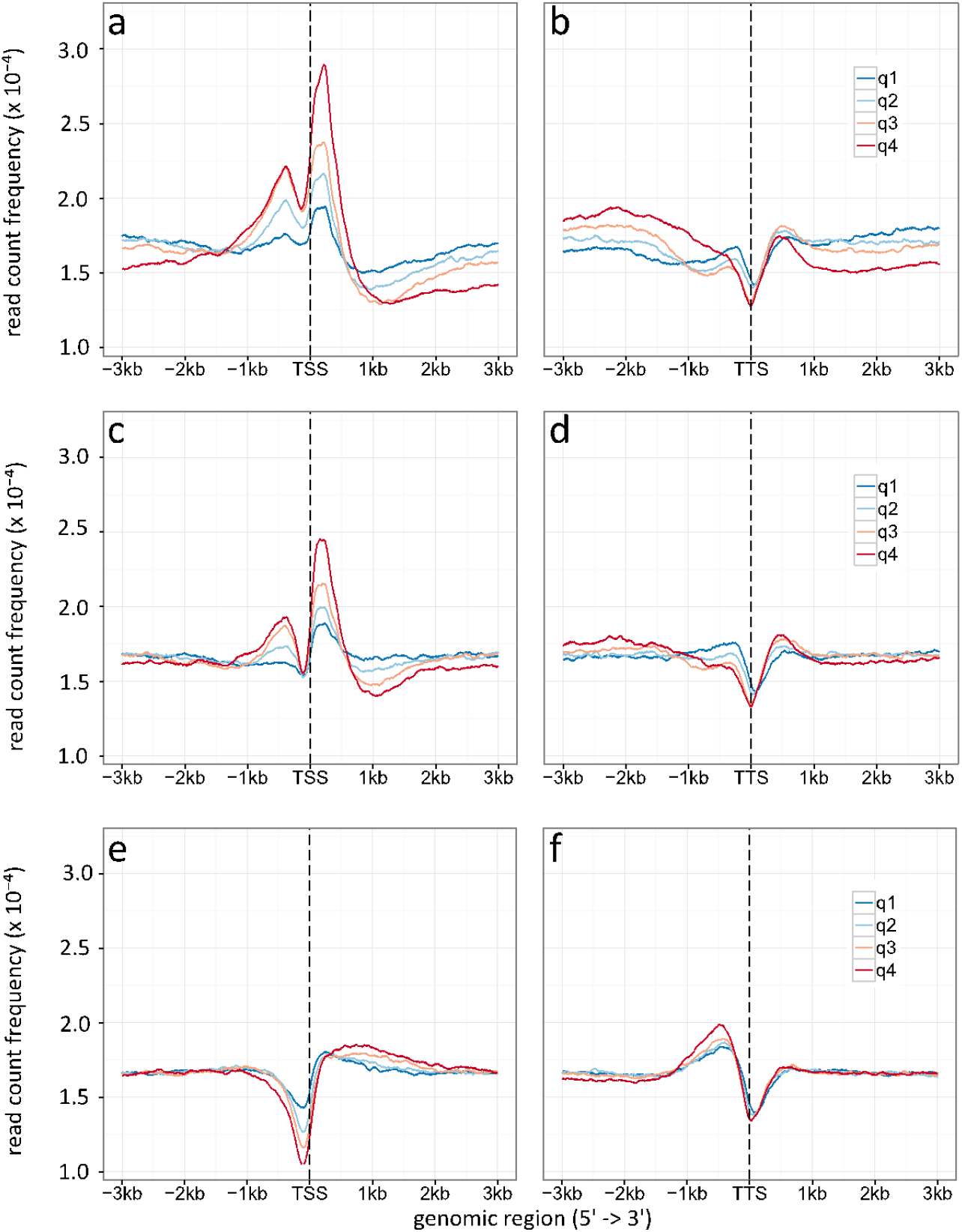
Density plot of TSS (a, c, e) and TTS (b, d, f) given for different gene expression levels (q1-q4). (**a, b**) Specific enrichment of H3K9ac, (**c, d**) H3K14ac and (**e, f**) H3 is given in read count frequency. Thereby q1 are the lower 25 %, q2 the medium lower 25 – 50 %, q3 are the medium high 50-75 %, q4 the higher 25 %.

**Figure S13.**
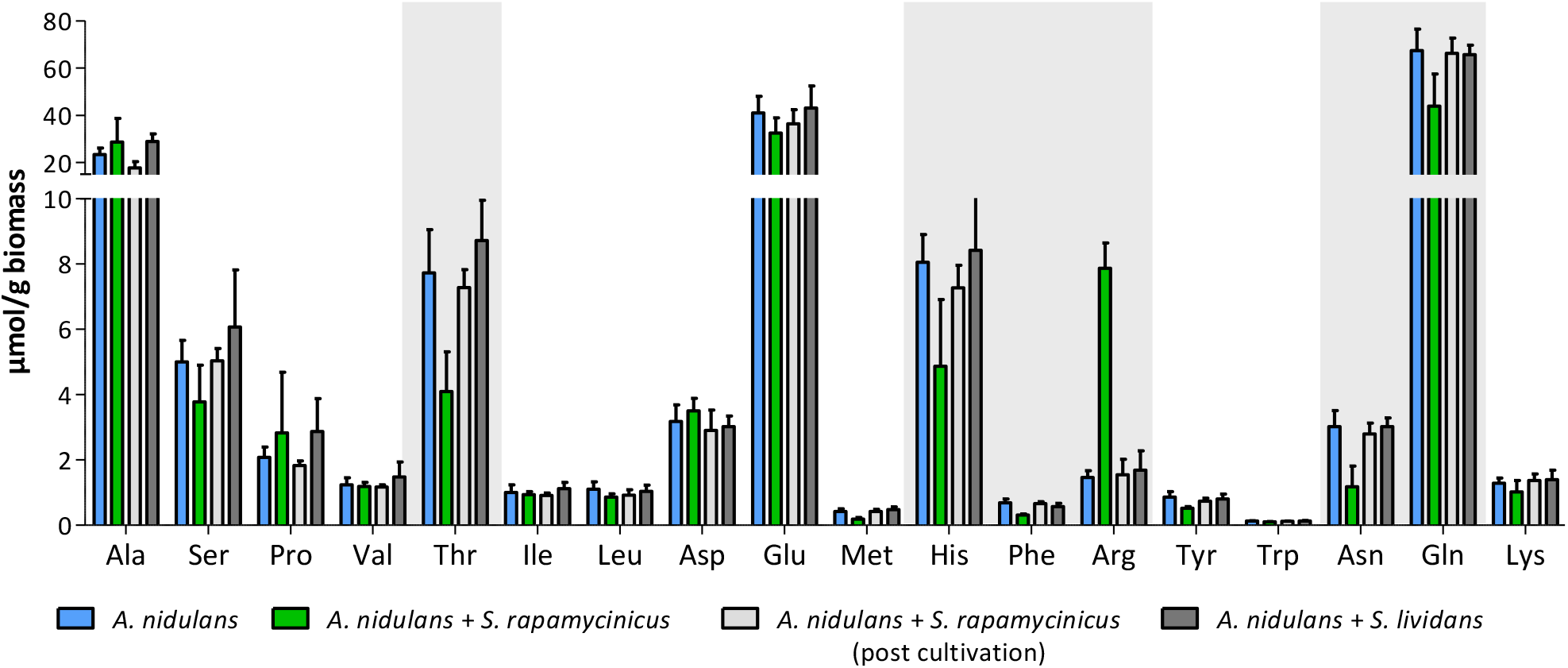
Intracellular amino acid concentration of *A. nidulans* in monoculture and co-culture with *S. rapamycinicus*. Co-cultivation with *S. lividans* and addition of *S. rapamycinicus* after 24 hours of cultivation served as negative controls. Furthermore, before extraction of amino acids the fungus (post cultivation) was also supplemented with *S. rapamycinicus* to exclude a bias resulting from bacterial amino acids. Threonine, histidine, phenylalanine, arginine, asparagine and glutamine showing different concentrations in co-culture compared to monoculture are highlighted in grey.

**Figure S14.**
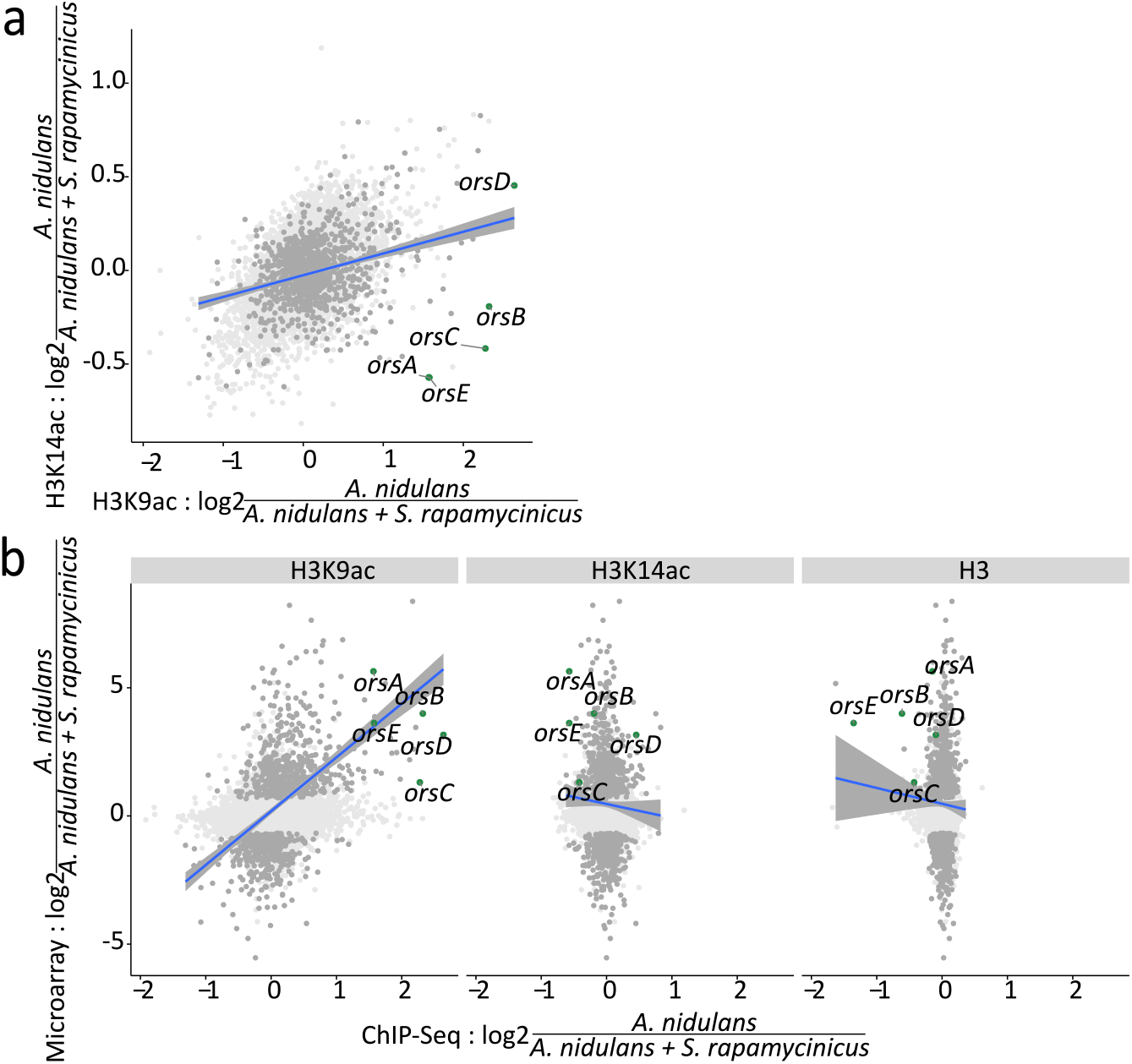
Relation between ChIP-seq and microarray data. The blue lines resemble the linear regression line based on the differentially expressed genes with an adjusted p-value of < 0.1 including the confidence interval shown in grey. (**a**) LFCs of H3K14ac plotted against LFCs of H3K9ac. Dots depicted in dark grey and green mark differentially expressed genes and *ors* cluster genes, respectively. (**b**) Pairwise comparison of LFCs of H3, H3K14ac and H3K9ac data with microarray data obtained during co-cultivation of *A. nidulans* with *S. rapamycinicus*.

## Supplementary tables

**Table S2.**
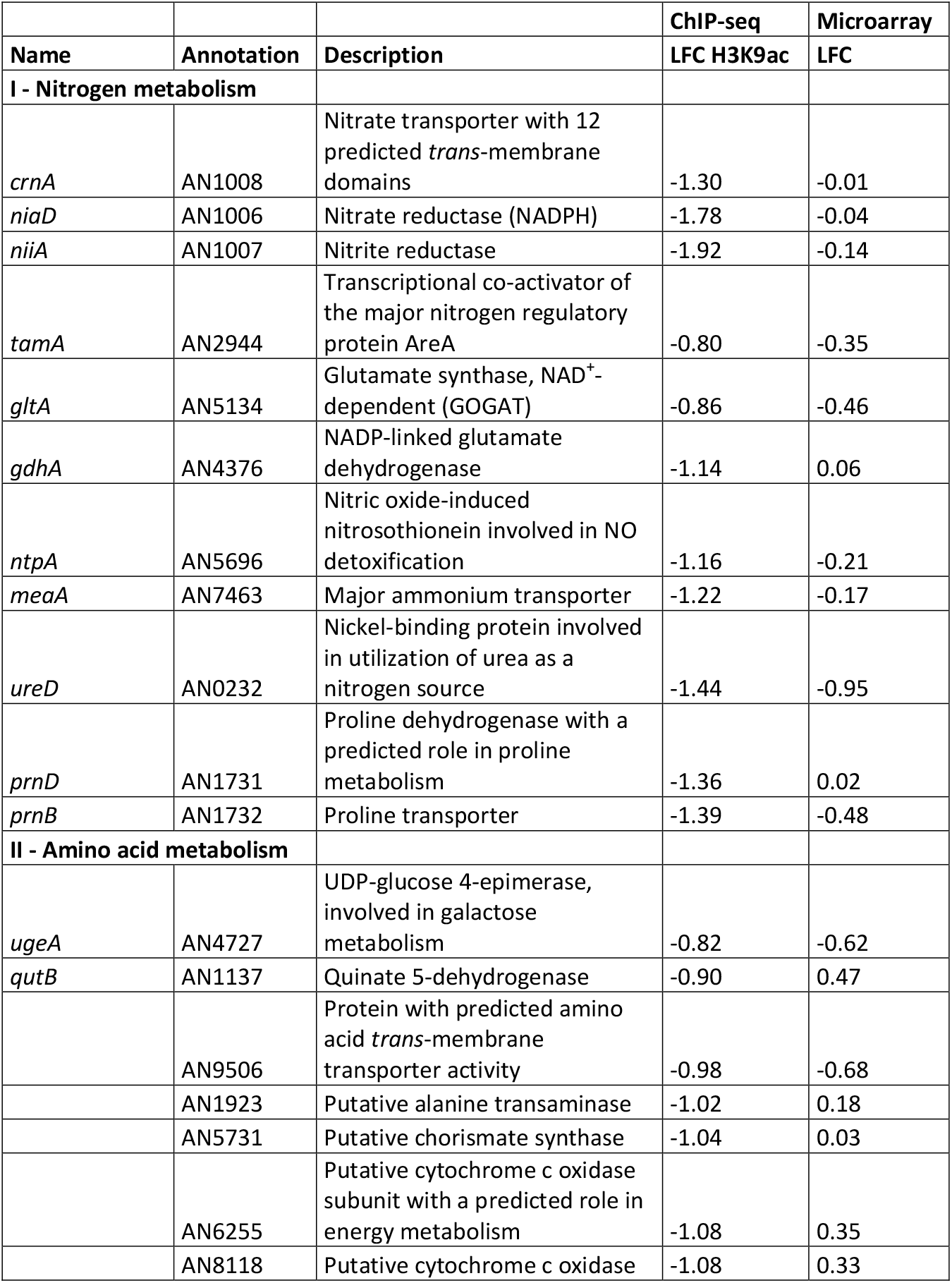
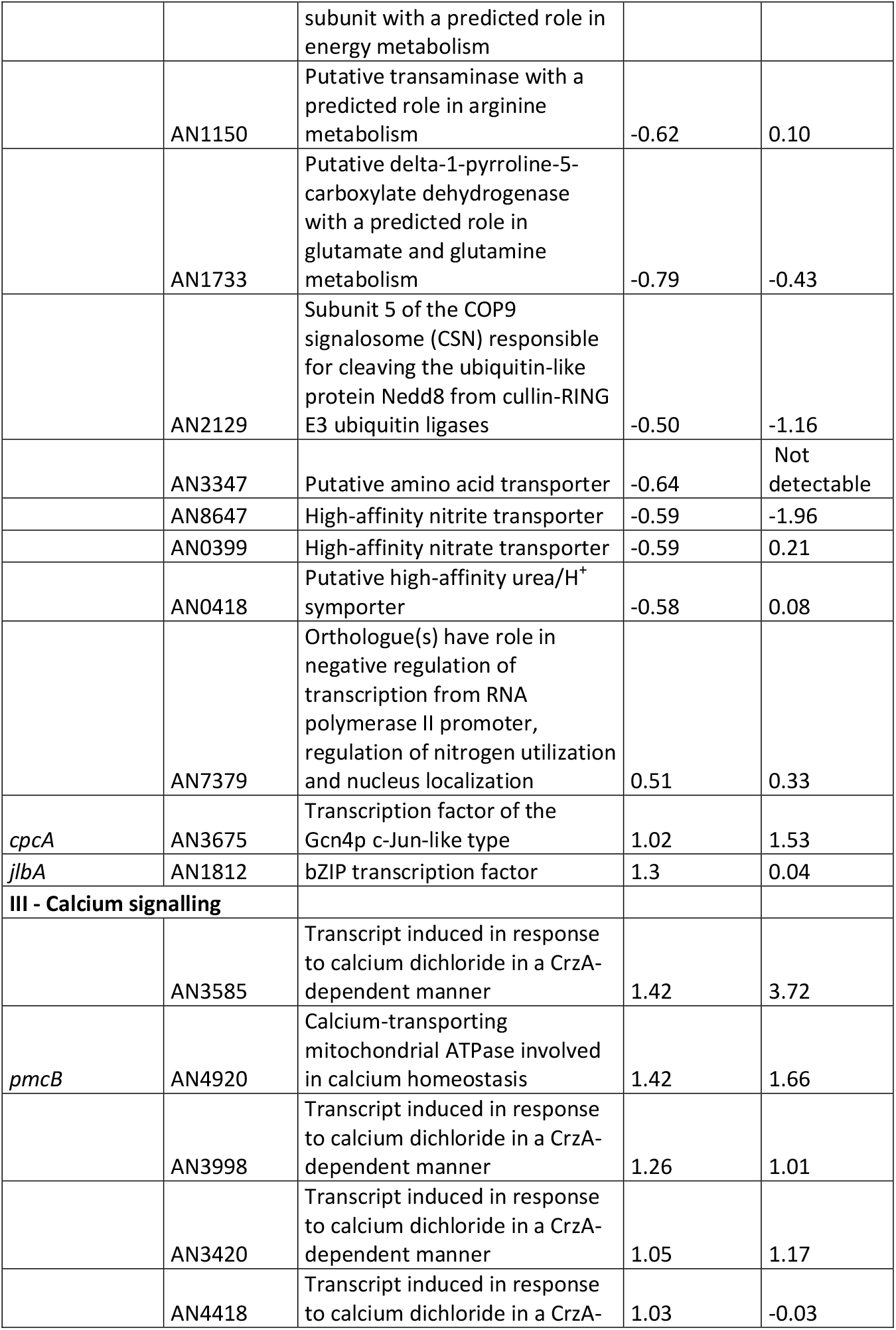
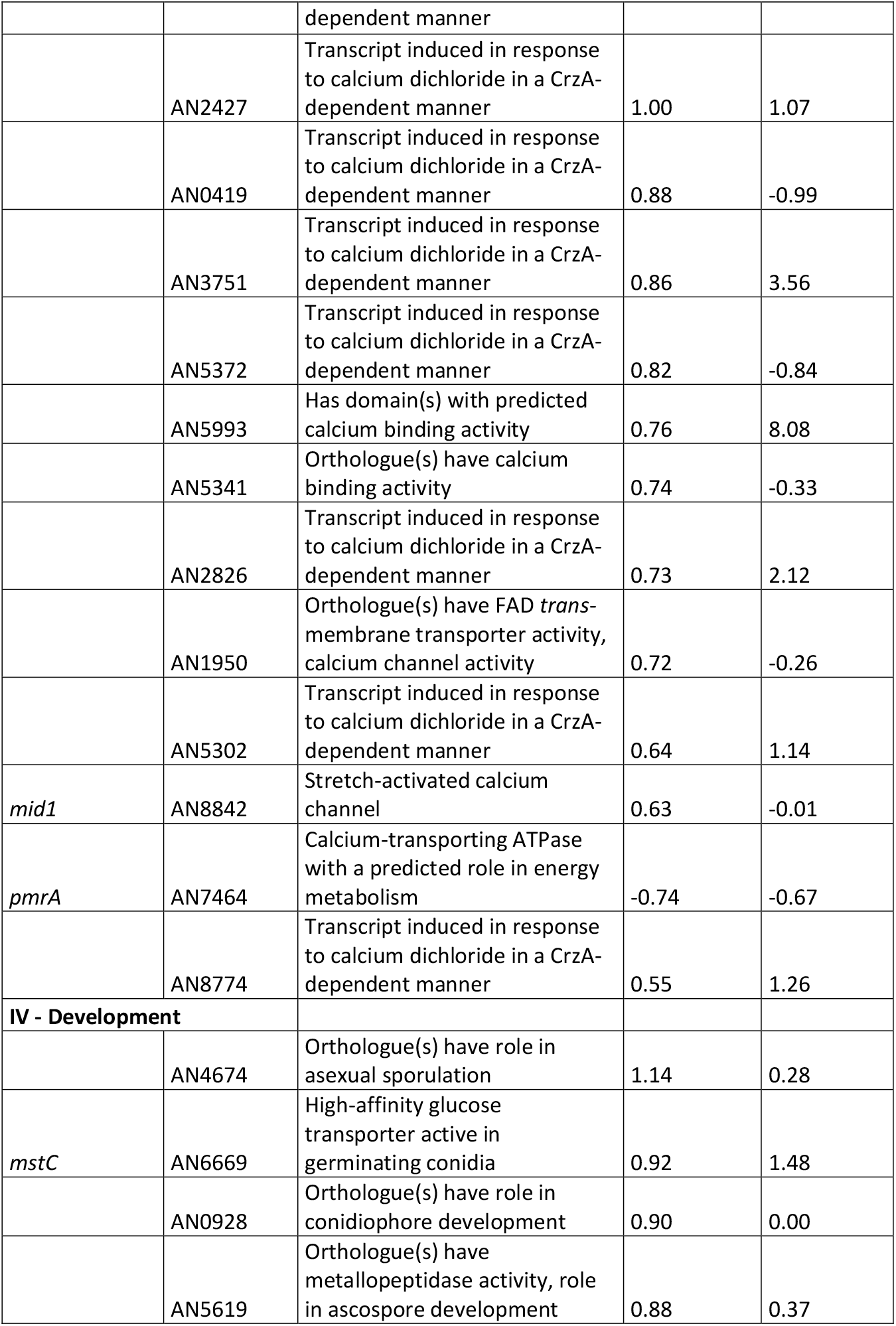
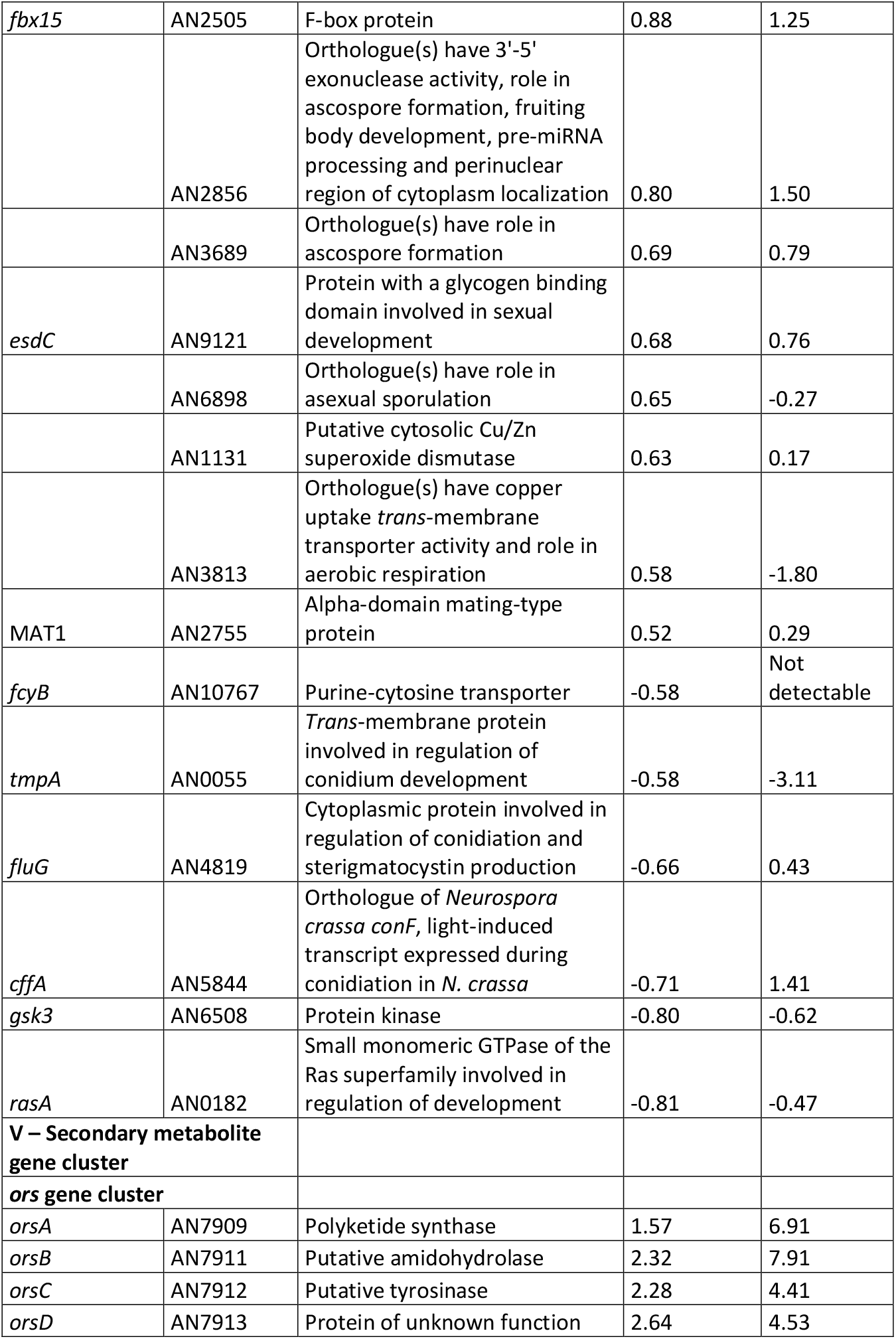
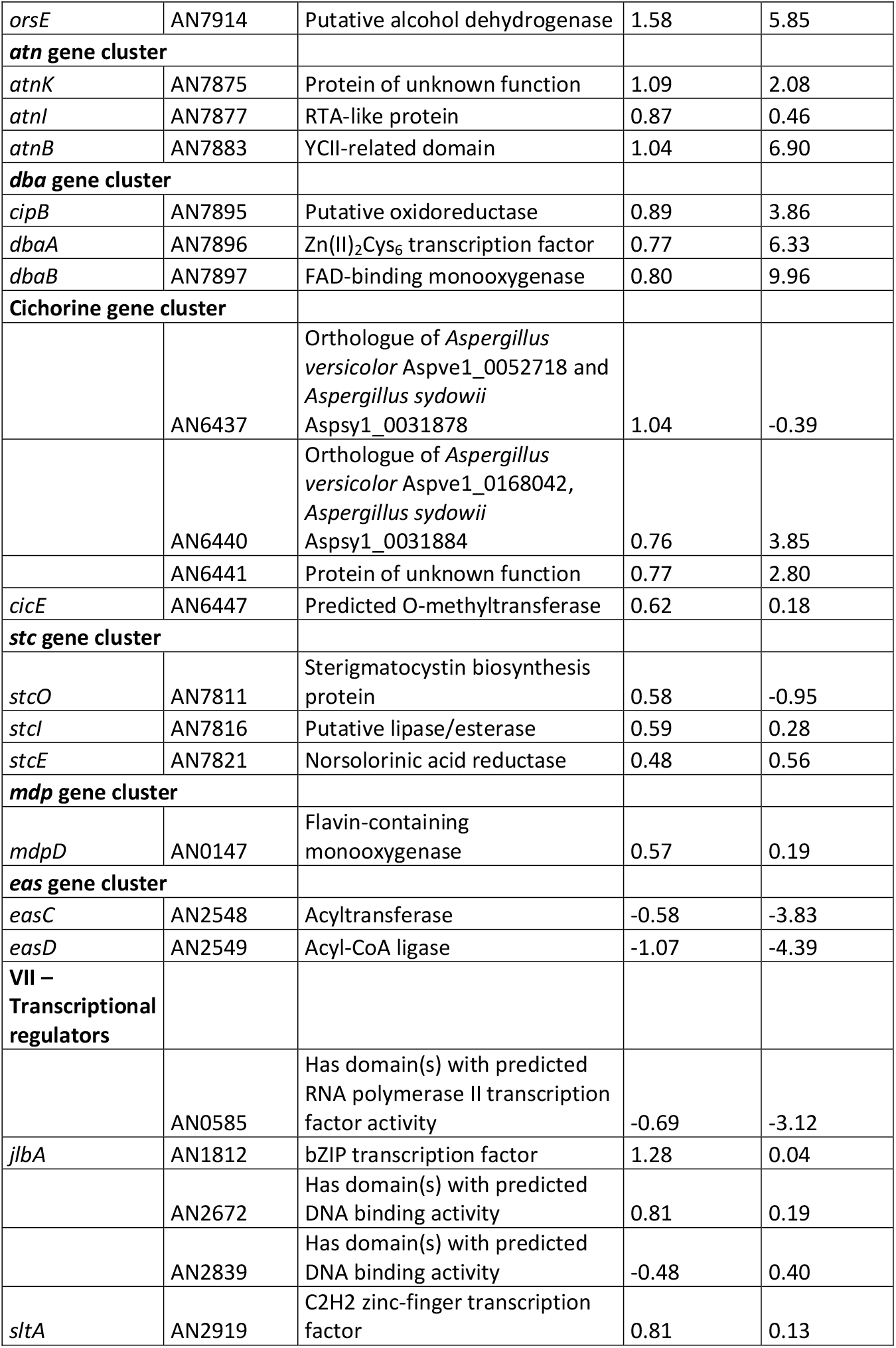
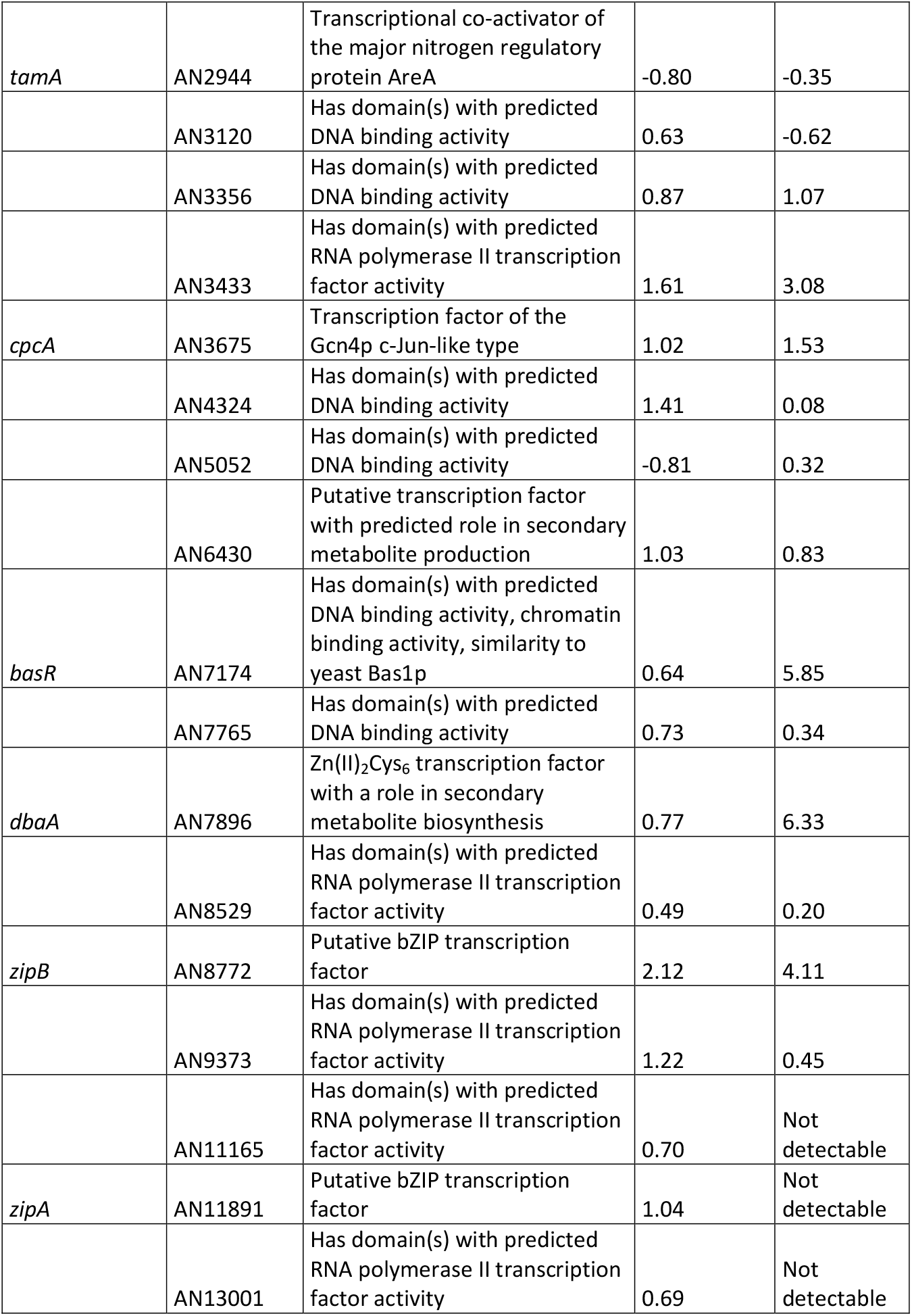
List of selected genes with differentially acetylated H3K9 and different expression.

**Table S4.**
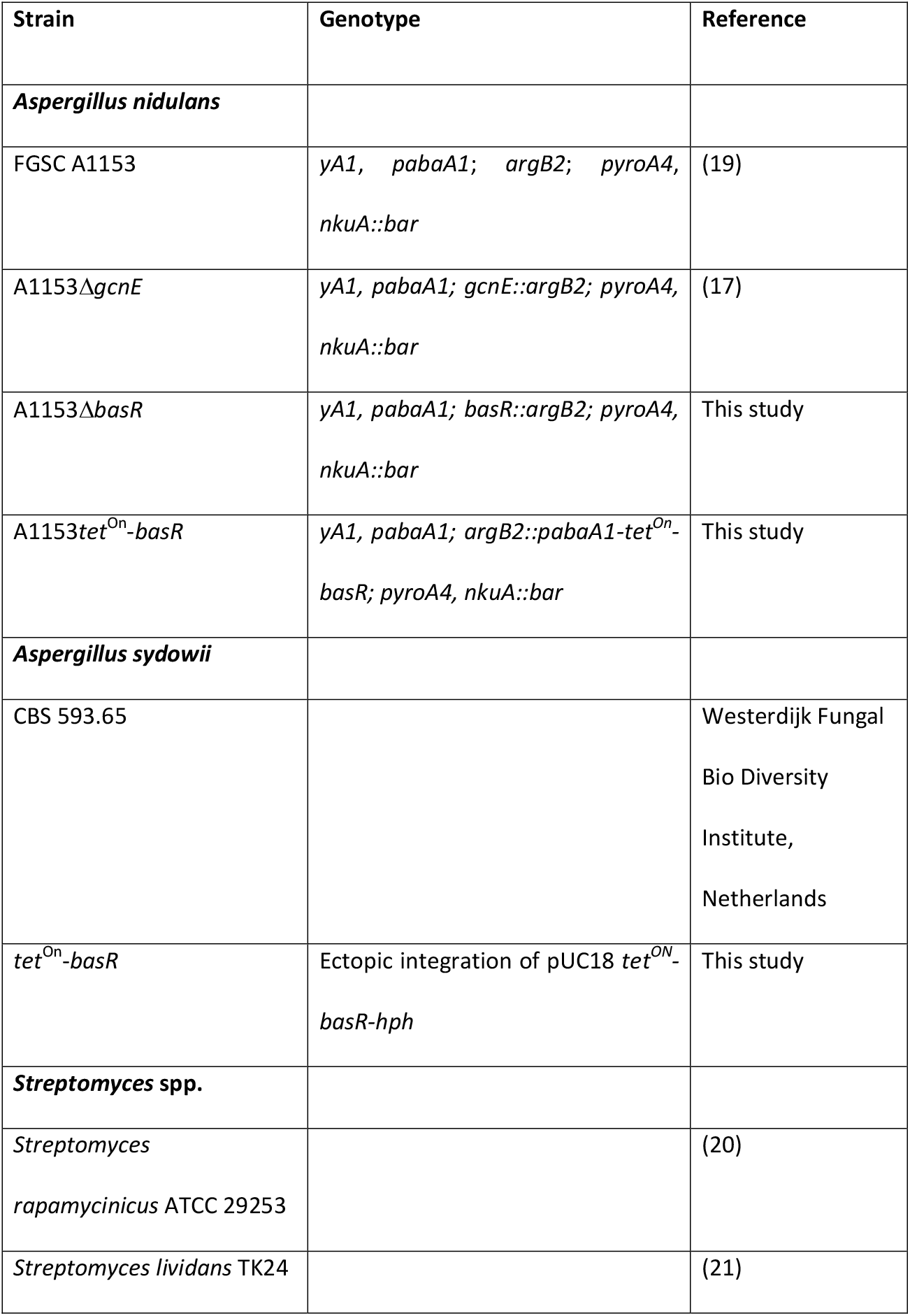
List of strains used in this study.

| Strain | Genotype | Reference |
| --- | --- | --- |
| ***Aspergillus nidulans*** |  |  |
| FGSC A1153 | *yA1, pabaA1; argB2; pyroA4, nkuA::bar* | (19) |
| A1153Δ*gcnE* | *yA1, pabaA1; gcnE::argB2; pyroA4, nkuA::bar* | (17) |
| A1153Δ*basR* | *yA1, pabaA1; basR::argB2; pyroA4, nkuA::bar* | This study |
| A1153*tet*<sup>On</sup>-*basR* | *yA1, pabaA1; argB2::pabaA1-tet*<sup>On</sup>-*basR; pyroA4, nkuA::bar* | This study |
| ***Aspergillus sydowii*** |  |  |
| CBS 593.65 |  | Westerdijk Fungal<br>Bio Diversity<br>Institute,<br>Netherlands |
| *tet*<sup>On</sup>-*basR* | Ectopic integration of pUC18 *tet*<sup>ON</sup>-*basR-hph* | This study |
| ***Streptomyces* spp.** |  |  |
| *Streptomyces rapamycinicus* ATCC 29253 |  | (20) |
| *Streptomyces lividans* TK24 |  | (21) |

**Table S5.**
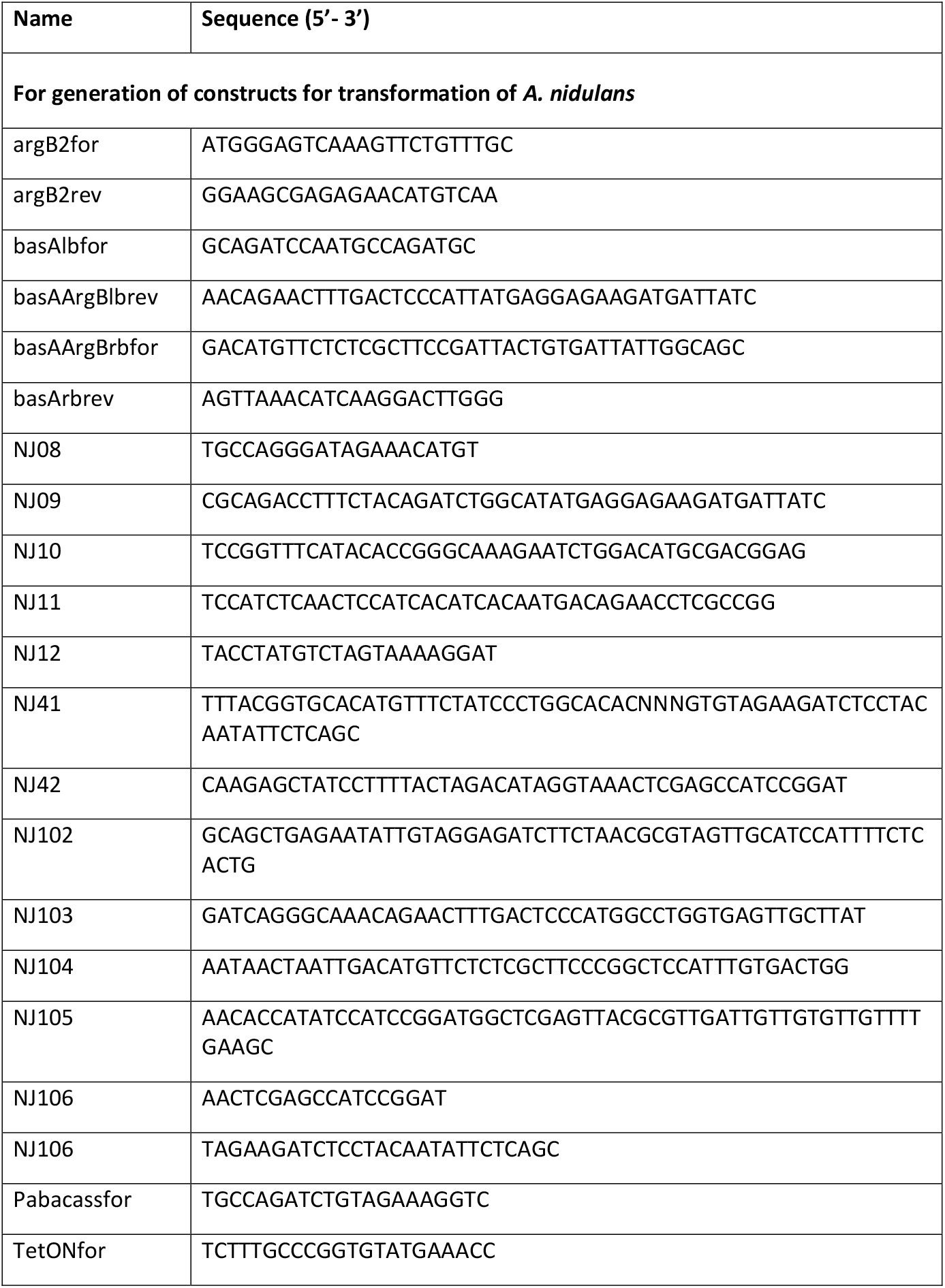
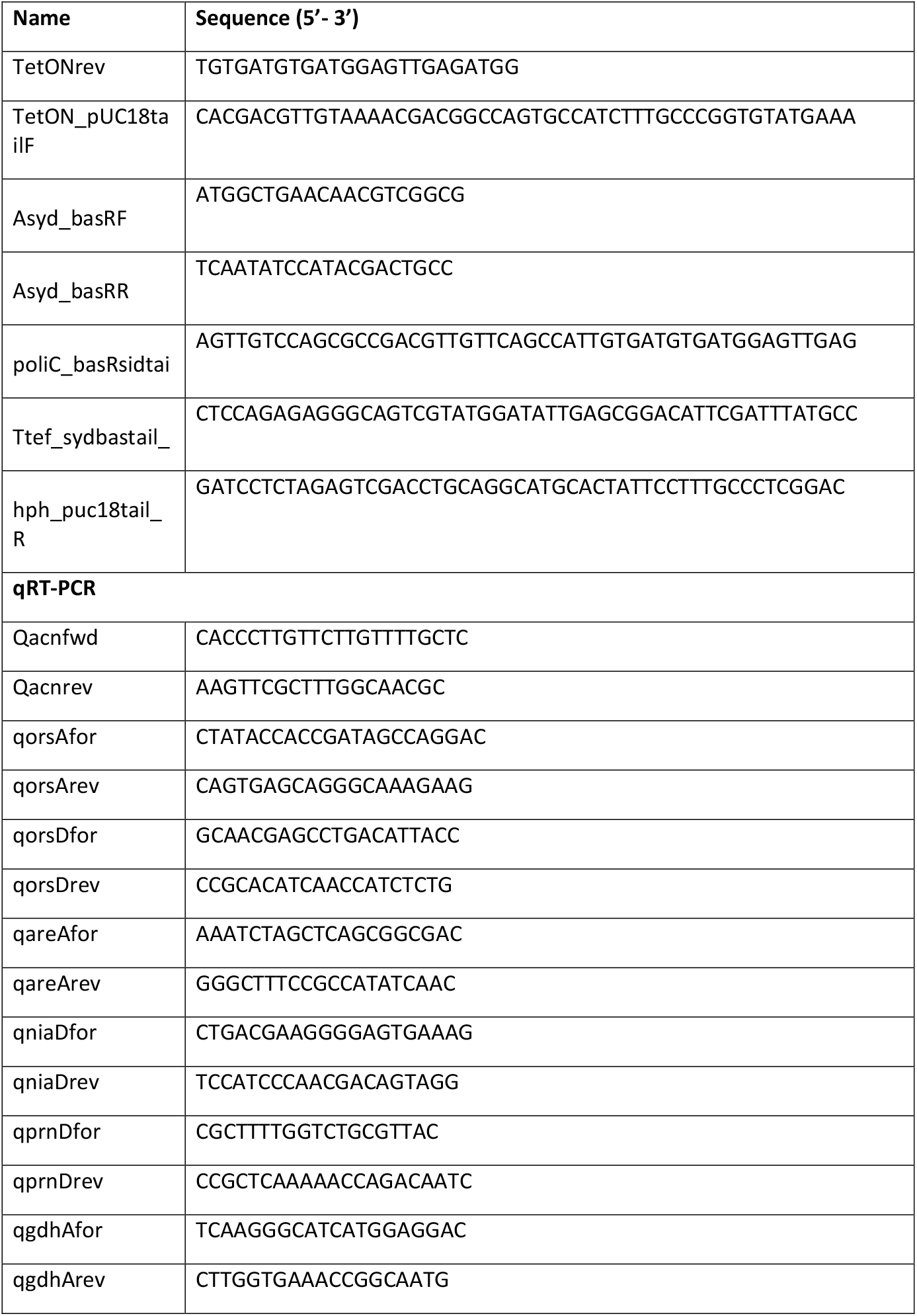
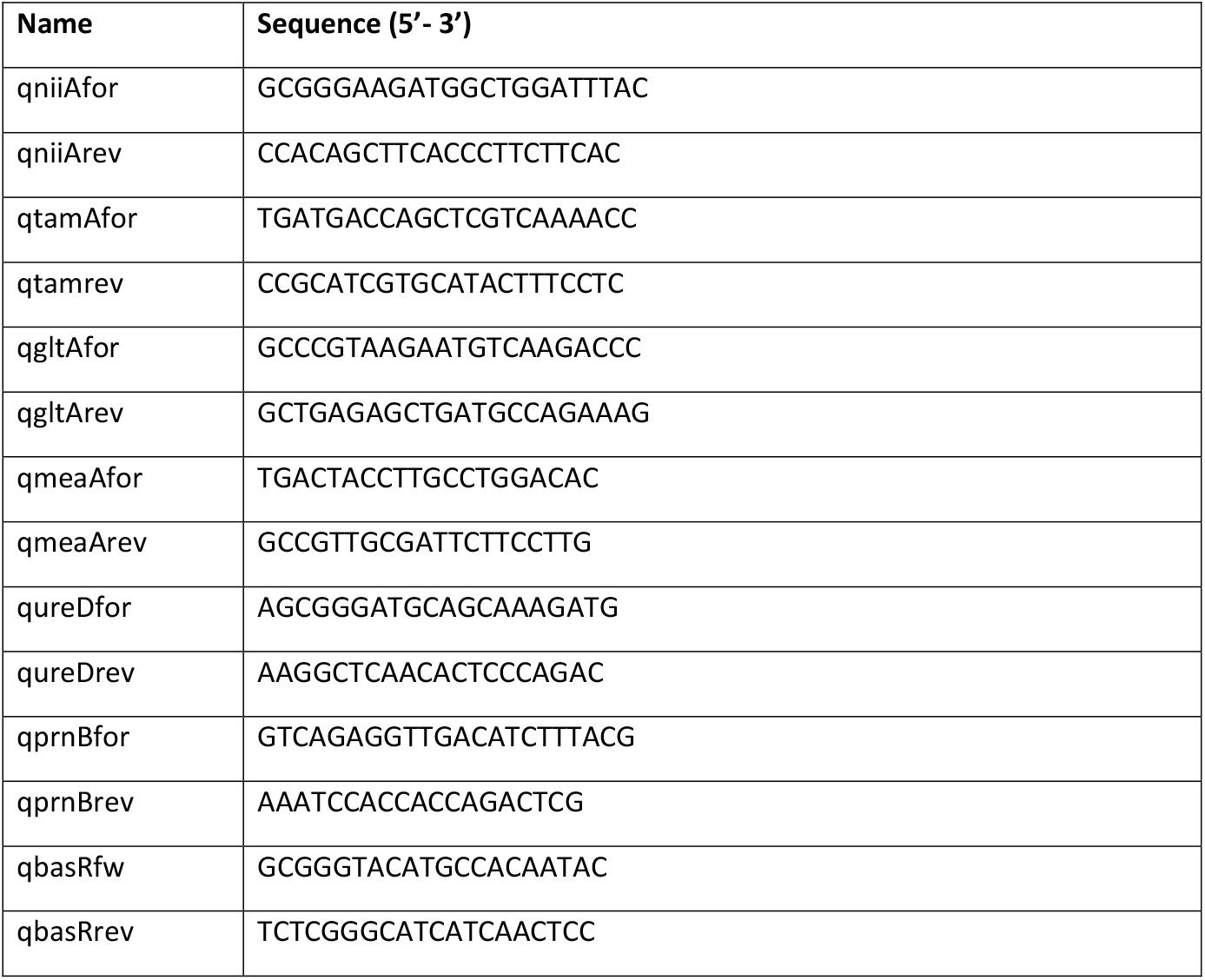
List of primers used in this study.

**Additional data table S1. (separate file)**

**Summary of DCS analysis of H3K9ac between *A. nidulans* monoculture and co-cultivation with *S. rapamycinicus*.** Significantly higher acetylated genes are marked in red and lower acetylated genes are marked in blue.

**Additional data table S3. (separate file)**

**Summary of ChIP-seq data**.

